# The BUBR1 pseudokinase domain promotes efficient kinetochore PP2A-B56 recruitment to regulate spindle checkpoint silencing and chromosome alignment

**DOI:** 10.1101/733378

**Authors:** Luciano Gama Braga, Angel F. Cisneros, Michelle Mathieu, Maxime Clerc, Pauline Garcia, Baptiste Lottin, Chantal Garand, Philippe Thebault, Christian R Landry, Sabine Elowe

## Abstract

The balance of phospho-signalling at outer-kinetochores during mitosis is critical for the accurate attachments between kinetochores and the mitotic spindle and timely exit from mitosis. In humans, a major player in determining this balance is the PP2A-B56 phosphatase which is recruited to the Kinase Attachment Regulatory Domain (KARD) of the Spindle Assembly Checkpoint protein Budding Uninhibited by Benzimidazole 1-related 1 (BUBR1) in a phospho-dependent manner. This event unleashes a rapid, switch-like phosphatase relay that reverses phosphorylation at the kinetochore, extinguishing the checkpoint and promoting anaphase entry. Here, we conclusively demonstrate that the pseudokinase domain of human BUBR1 lacks phosphotransfer activity and that it was maintained in vertebrates because it allosterically promotes KARD phosphorylation. Mutation or removal of this domain results in decreased PP2A-B56 recruitment to the outer kinetochore, attenuated checkpoint silencing and errors in chromosome alignment as a result of imbalance in Aurora B activity. We demonstrate that the functions of the BUBR1 pseudokinase and the BUB1 kinase domains are intertwined, providing an explanation for retention of the pseudokinase domain in certain eukaryotes.

## INTRODUCTION

Proper chromosome segregation during mitosis relies on the capacity of the spindle assembly checkpoint (SAC) to sense the attachment status of kinetochores and to delay progression to anaphase until unproductive attachment errors are corrected ^1, 2^. The SAC is an exquisitely conserved signalling cascade initiated by the recruitment of the kinase monopolar-spindle 1 (MPS1, also known as TTK) to kinetochores that are unattached to microtubules ^3–8^. MPS1 phosphorylation of the kinetochore scaffolding protein kinetochoreless-1 (KNL1) at a number of repetitive Met-Glu-Leu-Thr (MELT) motifs. Phosphorylated KNL1 then associates with the budding uninhibited by benzimidazole (BUB) proteins BUB3-BUB1 and subsequently recruits BUB3-BUBR1 (BUB1 Related-1) heterodimers in a pseudo-symmetrical manner ^11–16^. BUB1 phosphorylation also induces Mitotic Arrest Deficient 1 (MAD1) binding which supports the recruitment of MAD1-MAD2 heterodimers ^17–19^. MAD2, BUB3, and BUBR1, together form a potent and diffusible anaphase inhibitor, although the molecular details of the ultimate inhibitory complex remain unclear ^9, 20, 21^.

SAC silencing largely relies on timely recruitment of phosphatase activity to the outer-kinetochore to reverse early mitotic phosphorylation. Elegant studies have demonstrated that in vertebrates, recruitment of the PP2A phosphatase occurs mainly through direct interaction between one of several isoforms of the B56 adaptor subunit and the Kinetochore Alignment Regulatory Domain (KARD) of BUBR1 ^22–28^, although recent work has also suggested a role for B55-PP2A in SAC extinction ^29, 30^. BUBR1-bound B56-PP2A then promotes recruitment of PP1 to at least one of its receptors at the kinetochore, KNL1. Together, PP1 and PP2A drive a dephosphorylation relay which reverses early mitotic phosphorylation thereby permitting anaphase entry ^31–36^. BUBR1-bound PP2A has also been shown to destabilize the interaction between BUB1 and MAD1, and to reverse Aurora B phosphorylation events at the outer kinetochore to stabilize kinetochore-microtubule attachments ^35, 37^. Phosphorylation of the KARD by PLK1 at S676 and CDK1 at S670 enhances the binding of B56 to the core binding motif and is required in mammalian cells for optimal silencing of the SAC ^22, 24^. BUBR1 is thus in a unique position in the SAC cascade as one of the ultimate mediators of SAC signalling but also as one of the first players in its extinction.

Structure-function analysis of conserved motifs in BUBR1 orthologs in several model organisms has led to a remarkable understanding of this protein at the molecular level with the exception of the kinase domain. In most eukaryotes BUBR1 orthologues lack the kinase domain and are known as MAD3. BUB1 and BUBR1 are paralogs that are thought to have evolved from a single ancestral gene, termed MADBUB, present in the Last Eukaryotic Common Ancestor (LECA), through multiple gene duplication and parallel subfunctionalization events^38–40^. These events resulted in the retention of an active kinase domain across BUB1 orthologs, and its selective loss in MAD3 (Fig 1A). Remarkably, in a number of insects and vertebrates, the kinase domain was retained but its function remains controversial. A number of early studies suggested that characteristic BUBR1 mitotic hyperphosphorylation occurs through kinase activation and autophosphorylation resulting from a direct association of the kinase domain with the kinesin motor CENP-E ^41–43^. More recently, it has been proposed that vertebrate BUBR1 is in fact an unusual pseudokinase that has retained the catalytic triad, but is inactive as a result of the degeneration of the Gly-rich loop, together with an activation loop phosphorylation that is incompatible with catalysis ^38, 39, 44, 45^. Instead, the BUBR1 kinase domain has been shown to confer stability, and mutations in and around the C-terminal domain reduce total BUBR1 protein levels and are associated with the cancer predisposition disease Mosaic Variegated Aneuploidy (MVA) ^46–49^.

**Figure 1:**
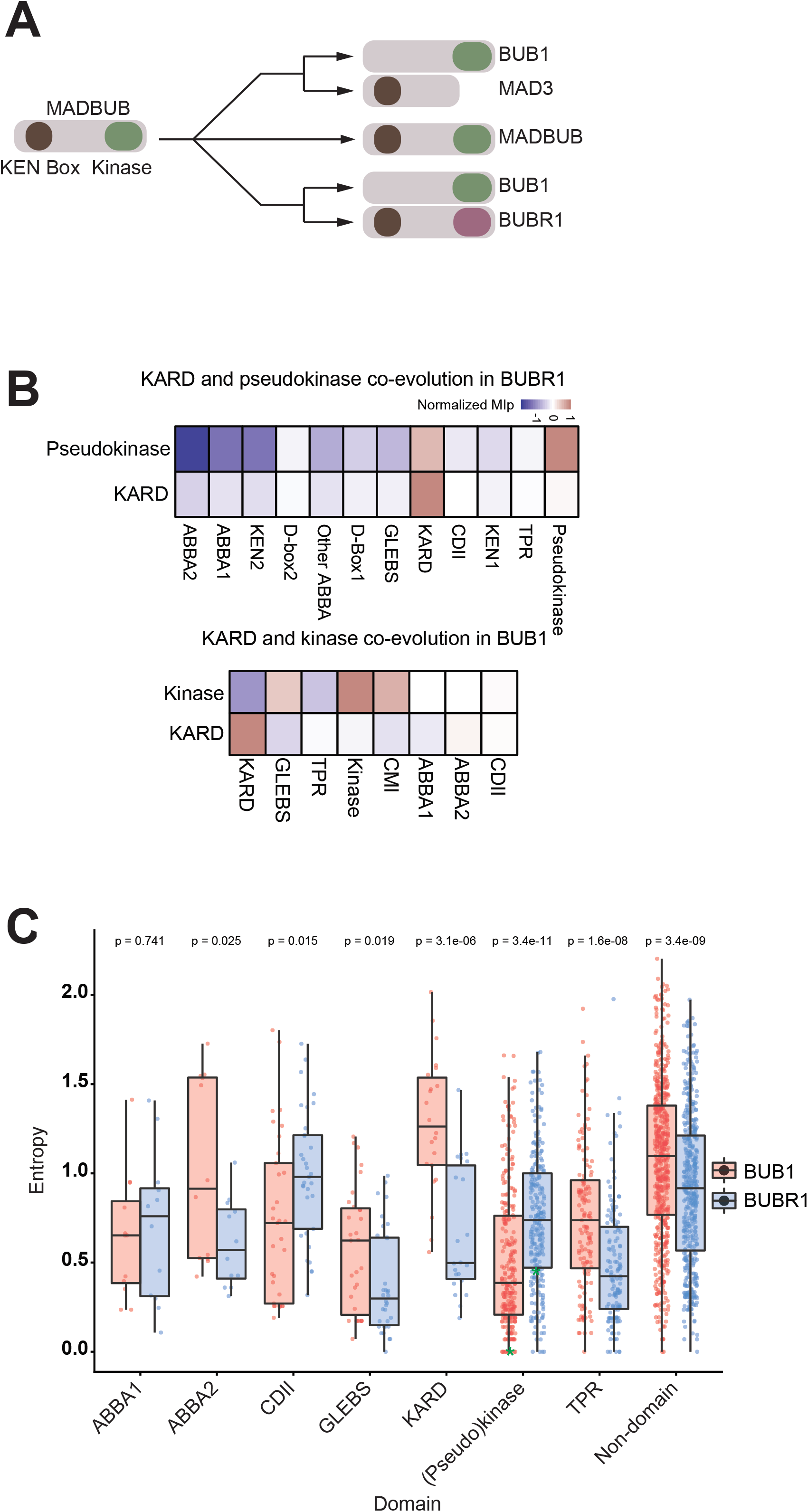
Co-evolution of the BUBR1 pseudokinase and KARD domains. A: Schematic representation of the evolutionary paths of the BUB1 and BUBR1-encoding genes. Their hypothetical LECA, a gene termed MADBUB, is believed to have presented a KEN box (in brown) and a kinase domain (in green), which are crucial for SAC activity. The hypothetical MADBUB ancestral protein has evolved in different configurations in animal and fungi. In the first pattern, the MADBUB gene retains both the KEN box and kinase domain. In the second scenario, the kinase is truncated in one paralog, whereas the other gene retains the kinase but loses the KEN box as is seen in certain yeasts and nematodes. In most vertebrates a third scenario occurred where the KEN box-containing gene retained a kinase domain lacking catalytic function (domain in purple). Some exceptions to this exist and are discussed in detail in Suijkerbuijk et al. ^38^ and Vleugel et al. ^40^ and Tromer et al. ^39^. B: Co-evolution heatmap of BUBR1 and BUB1 KARD and (pseudo)kinase domains. The normalized mutual information of a protein (MIp) directly correlates to co-evolving positions in protein families. C: Entropy comparison of domains from BUB1 and BUBR1 protein families. Entropy inversely correlates to the level of conservation of a domain. Each point corresponds to an aligned residue in its corresponding domain. Residues equivalent to catalytic asperates (D917 in hBUB1, D882 in hBUBR1) are marked with a green asterisk.

The relative stability of BUBR1 lacking the C-terminus ^49^ together with loss of the kinase domain in most eukaryotes prompted us to explore whether this domain confers other as yet uncharacterized attributes to BUBR1 in species where it has been retained, using human BUBR1 as a model. Here we demonstrate that BUBR1 mitotic hyperphosphorylation is not a result of autoactivation and autophosphorylation, and confirm previous results that BUBR1 is a pseudokinase incapable of phosphotransfer activity. We demonstrate that mutations in the pseudokinase domain not only destabilize BUBR1 but also significantly attenuate phosphorylation in the neighbouring KARD at S670 and S676. Consequently, mutation or removal of the pseudokinase domain results in decreased PP2A-B56 recruitment to the outer kinetochore, attenuated SAC silencing and errors in chromosome bi-orientation as a result of imbalance in Aurora B activity. We conclude that the pseudokinase domain functions as an allosteric regulator of KARD phosphorylation in a manner that requires the BUB1 kinase domain, providing an explanation for retention of the pseudokinase domain.

## RESULTS

### Co-evolution of the KARD and pseudokinase domains of BUBR1

In an attempt to identify potential functions for the BUBR1 pseudokinase domain, we sought to determine whether there is a co-evolutionary relationship between the pseudokinase and additional domains of BUBR1. To do this, we analyzed sequences from Ensembl Orthologs as well as previously collated sequences ^39, 50^. We measured co-evolution between domains for both BUB1 and BUBR1 by looking at the mutual information between residues and applying normalizations to correct for background noise ^51^ and intrinsic differences in variability between different domains (see methods). The resulting metric, normalized MIp, is positive when two domains co-evolve more strongly with each other than expected from background signal. Independently, we looked at the co-presence of domains using Pearson correlation as reported by Tromer et al ^39, 51^. Both approaches suggest that the pseudokinase domain of BUBR1 co-evolves with the KARD in terms of both sequence variation and domain co-presence (and Fig 1B, **Suppl. Fig 1A, B**). KARD-like motifs can also be recognized in BUB1, and they show a certain similarity to BUBR1 KARD sequences (^39^ **Supp. Fig 1C**). However, although the core B56 binding motif is easily identified in hBUB1, it was non-functional, likely as a result of the absence of C-terminal acidic residues (**Supp. Fig 1D** ^22^). In agreement, both co-evolution and Pearson correlation were weaker between the kinase domain and KARD-like sequences of BUB1 than for BUBR1 (Fig 1B, **Suppl. Fig 1 E, F).**

The intriguing evolutionary path of the BUB1 and BUBR1/MAD3 genes also prompted us to analyze the degeneration of their various shared domains in an attempt to understand how the kinase domain and KARD evolved in both paralogs. Diversity as measured by Shannon entropy showed that, while the BUB1 kinase, the BUBR1 pseudokinase, and BUBR1 KARD are relatively conserved, the BUB1 KARD-like domain degenerates more rapidly (Fig 1C). This suggests that co-functionality between the KARD and pseudokinase domain of BUBR1. Moreover, mapping evolution of the catalytic aspertate in both BUB1 and BUBR1 demonstrated divergence in BUBR1 compared to BUB1, in agreement with lack of BUBR1 catalytic activity in most orthologs (see position marked with asterisk in Fig 1C).

### The pseudokinase domain is essential for BUBR1 stability and phosphorylation

BUBR1 hyperphosphorylation is a hallmark of mitosis, resulting in a characteristic double-band pattern in Western blots, but the origin of this hyperphosphorylation remains actively disputed. We and others have previously shown that this is PLK1-dependent ^52–57^. However, it has also been suggested that this hyperphosphorylation is a result of PLK1 activation of BUBR1 kinase activity ^54, 56^, and independently, of CENP-E binding and BUBR1 autophosphorylation in a checkpoint-dependent manner ^41–43^. These conclusions, however, are not compatible with observations that BUBR1 is catalytically inactive ^38, 44, 45^. To address these discrepancies, we generated a panel of BUBR1 pseudokinase domain mutants and asked whether they become hyperphosphorylated during mitosis. In HeLa S3 cells treated with nocodazole, mutation of two residues in the remnants of the ancestral catalytic triad, K795R (single kinase dead, SKD) and D911A, individually or in tandem (double kinase dead, DKD) highly destabilized BUBR1, in agreement with previous work ^38, 49^, but also significantly attenuated BUBR1 hyperphosphorylation as measured by retarded gel mobility (Fig 2A). When expression levels were normalized and activity tested by *in vitro* kinase assays, no appreciable autophosphorylation or phosphorylation of exogenous substrates was observed for BUBR1, although it was readily detected for BUB1, demonstrating that hyperphosphorylation is independent of putative BUBR1 kinase activity (**Supp. Fig 2A**). To determine whether BUBR1 hyperphosphorylation correlates with protein stability, we tested the gel mobility shift of BUBR1 MVA mutants that occur at or around the pseudokinase domain, and which are known to impair protein stability ^49^. We find that MVA mutants R727C, R814H, L844F, and L1012P were even less phosphorylated than K795 mutants, again in line with the idea that BUBR1 stability and hyperphosphorylation, but not catalytic activity, are correlated (Fig 2B).

**Figure 2:**
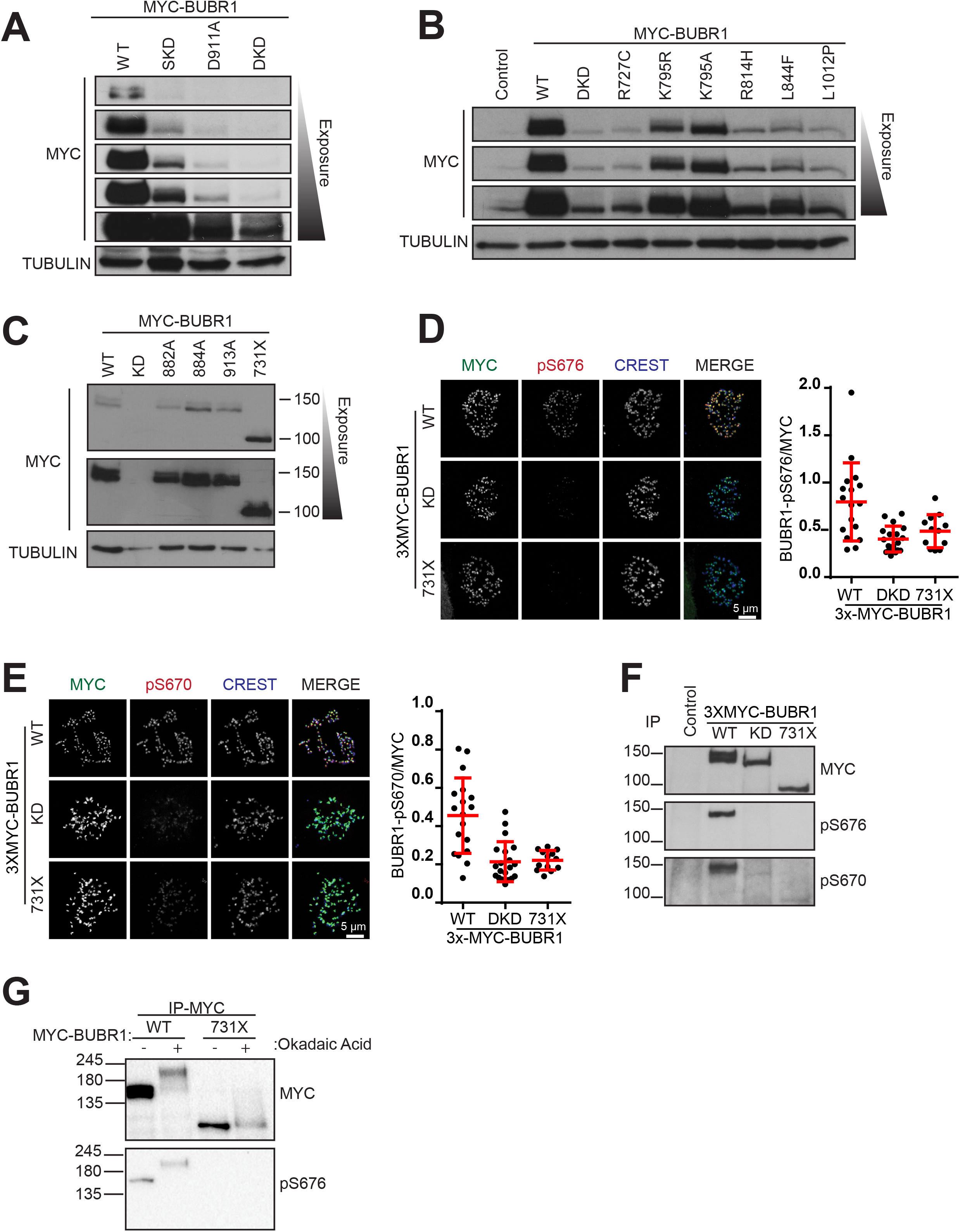
BUBR1 KARD Phosphorylation is attenuated in BUBR1 pseudokinase domain mutants. A-C: MYC (upper panel) and tubulin (lower panel) Immunoblots of lysates from nocodazole-arrested HeLaS3 cells transfected with the indicated 3xMYC-tagged BUBR1 mutants. Multiple MYC exposures are shown. D and E: HeLa cells stably expressing 3XMYC-BUBR1-WT, DKD, and 731X were depleted of endogenous BUBR1 by siRNA and treated with nocodazole. Fixed cells were stained with anti-MYC (green), anti-CREST (blue) and anti-BUBR1-pS676 or anti-BUBR1-pS670 (red). The graph indicated quantitation of phosphosignal intensity relative to MYC. F: MYC-BUBR1 was immunoprecipiated from stable cell lines arrested in mitosis by nocodazole treatment and immunoblotted with anti-pS676 and anti-pS670 antibodies. G: MYC-BUBR1 was immunoprecipitated from stable cell lines arrested in Nocodazole (1 µg/mL) overnight and treated with Okadaic Acid for 1 hour before lysis. Lysates were immunoblotted with anti-pS676 antibodies.

To definitively demonstrate that BUBR1 is a genuine pseudokinase, we next inquired if kinase mutations that interfere with catalysis of active kinases could influence BUBR1 hyperphosphorylation without compromising protein stability. We found that mutation of the putative catalytic aspartate of BUBR1 D882A (HR*D* in motif VI) did not disrupt protein integrity. Importantly, when expressed in nocodazole-arrested HeLaS3 cells, BUBR1-D882A displayed normal hyperphosphorylation, confirming that the gel mobility shift of BUBR1 is not due to autophosphorylation (Fig 2C). We also identified two other stable BUBR1 mutants that support this conclusion; S884A (catalytic loop motif VI) and S913A in motif VII (DF*S* motif, cation binding loop), both of which conform to PLK1 consensus sites. Considering that PLK1 has been implicated in BUBR1 phosphorylation and activation ^56^, and the significance of these motifs in functional kinase domains, we reasoned that these sites may contribute to any potential catalytic activity. However, both S884A and S913A mutations exhibited protein stability and mitotic hyperphosphorylation comparable to wild-type (WT) BUBR1. In line with the idea that optimal conformational stability regulates BUBR1 phosphorylation, a BUBR1 mutant that entirely lacks the pseudokinase domain (BUBR1-731X) but renders the protein hyperstable relative to the WT ^49^, also did not display a gel mobility shift (Fig 2C).

To explore the effect of the pseudokinase on BUBR1 phosphorylation in more detail, we monitored KARD phosphorylation at S676 and S670 with phospho-specific antibodies ^52, 53^. In agreement with previous reports, both residues were highly phosphorylated in prometaphase-arrested cells depleted of endogenous BUBR1 but expressing BUBR1-WT. However, in cells expressing the hyperstable BUBR1-731X or the unstable BUBR1-DKD, phosphorylation at both KARD sites was reduced (Fig 2D, E; **Suppl. Fig 2B-C**). Phosphospecific signals were normalized by total BUBR1 levels, indicating that this is not an effect of variability in BUBR1 kinetochore levels. The same results were obtained by Western blotting (Fig 2F). To determine whether BUBR1 pseudokinase mutants are at all amenable to phosphorylation, we treated cells expressing BUBR1-WT and -731X with okadaic acid to inhibit endogenous phosphatases which resulted in a mobility upshift in both instances but no restoration of S676 phosphorylation in BUBR1-731X suggesting that the pseudokinase is absolutely required for proper phosphorylation of the KARD (Fig 2G). Taken together, our data confirm that hBUBR1 is indeed a catalytically inactive during mitosis, and that the presence of a stable pseudokinase domain *per se* and not its catalytic activity is a requisite for hBUBR1 mitotic phosphorylation in the KARD.

### The BUBR1 pseudokinase domain is required for PP2A-B56 kinetochore recruitment and downstream phosphatase signalling

PP2A-B56 binding to the outer kinetochore is mediated through the KARD of BUBR1 and is strongly promoted by phosphorylation at the aforementioned PLK1 and CDK1 sites ^22, 24^. Therefore, the decrease in KARD phosphorylation caused by pseudokinase domain truncation or instability prompted us to assess if PP2A-B56 binding to BUBR1 is attenuated in pseudokinase mutants. To ensure homogenous expression of BUBR1 mutants, we generated inducible stable cell lines expressing 3xMYC-GFP-tagged BUBR1-WT, -SKD, -DKD, and -731X and titrated doxycycline to obtain equivalent kinetochore levels of the different BUBR1 constructs (**Suppl. Fig 3A**). We analyzed HA-B56α kinetochore recruitment in nocodazole-arrested cells depleted of all B56 isoforms by siRNA ^32^. As expected, B56α localized at kinetochores of prometaphase arrested cells, but it was greatly reduced in BUBR1 depleted cells (**Suppl. Fig 3B**). Expression of BUBR1-WT in cells depleted of endogenous BUBR1 restored kinetochore localization of B56α, whereas expression of BUBR1-731X, -DKD or -SKD at similar levels did not (Fig. 3A). Therefore, the BUBR1 pseudokinase domain regulates PP2A-B56 docking to the outer kinetochore, likely through KARD phosphorylation.

**Figure 3:**
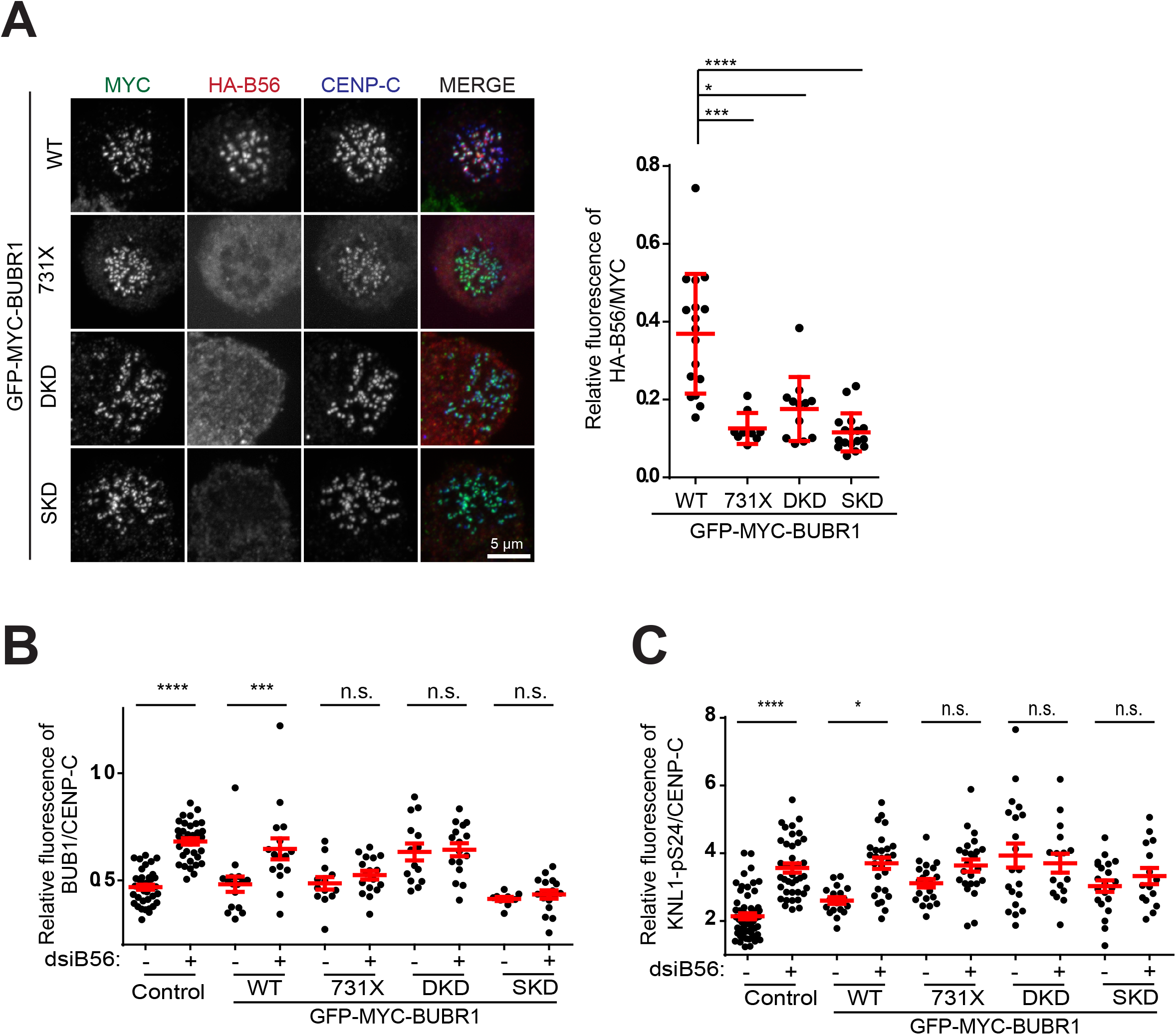
Signalling events downstream of KARD phosphorylation are attenuated in BUBR1 pseudokinase domain mutants. A: Effect of pseudokinase mutation on B56 localization in stable cell lines expressing GFP-MYC-BUBR1-WT, DKD, SKD, or 731X at equivalent levels. Cells expressing the indicated BUBR1 plasmids were depleted of endogenous BUBR1 and all B56 isoforms by dsiRNA treatment and transfected with HA-B56α, followed by nocodazole arrest overnight. Fixed cells were stained with anti-HA, anti-MYC, and anti-CENP-C antibodies. The graph indicates relative kinetochore levels of HA-B56α. B-C: Quantification of the relative BUB1 (B) and KNL1-pS24 (C) signals at kinetochores in the presence or absence of endogenous B56. Cells expressing similar levels of BUBR1 pseudokinase mutants were depleted of all endogenous B56 isoforms and BUBR1, and fixed for immunofluorescence with anti-BUB1 (B) or anti-KNL1-pSer24 (C), anti-MYC, and anti-CENP-C.

The BUBR1-bound pool of PP2A-B56 is responsible for dephosphorylating KNL1, which induces SAC silencing by two major mechanisms: first, by dephosphorylating PP1 phosphatase recruitment motifs on KNL1 (including S24 in the SILK motif, ^58, 59^) promoting its binding and further dephosphorylation of the kinetochore and secondly, by targeting KNL1 MELT motifs and inducing BUB1, BUB3 and BUBR1 delocalization ^31, 35, 37^. We reasoned that cells expressing BUBR1 pseudokinase mutants would exhibit comparatively weaker changes in these phosphosites after B56 depletion as a result of loss of PP2A-B56 signalling. In both control and BUBR1-WT expressing cells, we observed a significant increase in BUB1 localization to kinetochores, and in KNL1 S24 phosphorylation, as expected (Fig 3B-C). However, expression of BUBR1-731X, -DKD, or -SKD resulted in smaller changes that were not statistically significant. These observations suggest that the pseudokinase domain can indeed modulate downstream signalling from PP2A-B56.

### BUBR1 pseudokinase mutants undermine PP2A-B56 functions during mitosis

Considering the role of KARD phosphorylation in B56 interaction, we next reasoned that SAC silencing may be weakened in cells expressing BUBR1 pseudokinase mutants that display compromised PP2A-B56 kinetochore localization. To test this, we first arrested cells in prometaphase with nocodazole and induced mitotic exit by the addition of the Mps1 inhibitor reversine. Control cells, and cells expressing BUBR1-WT exited mitosis with similar kinetics. Strikingly, BUBR1-731X, -DKD and -SKD all caused a delay in mitotic exit, indicating that the pseudokinase domain is required for efficient SAC silencing (Fig 4A).

**Figure 4:**
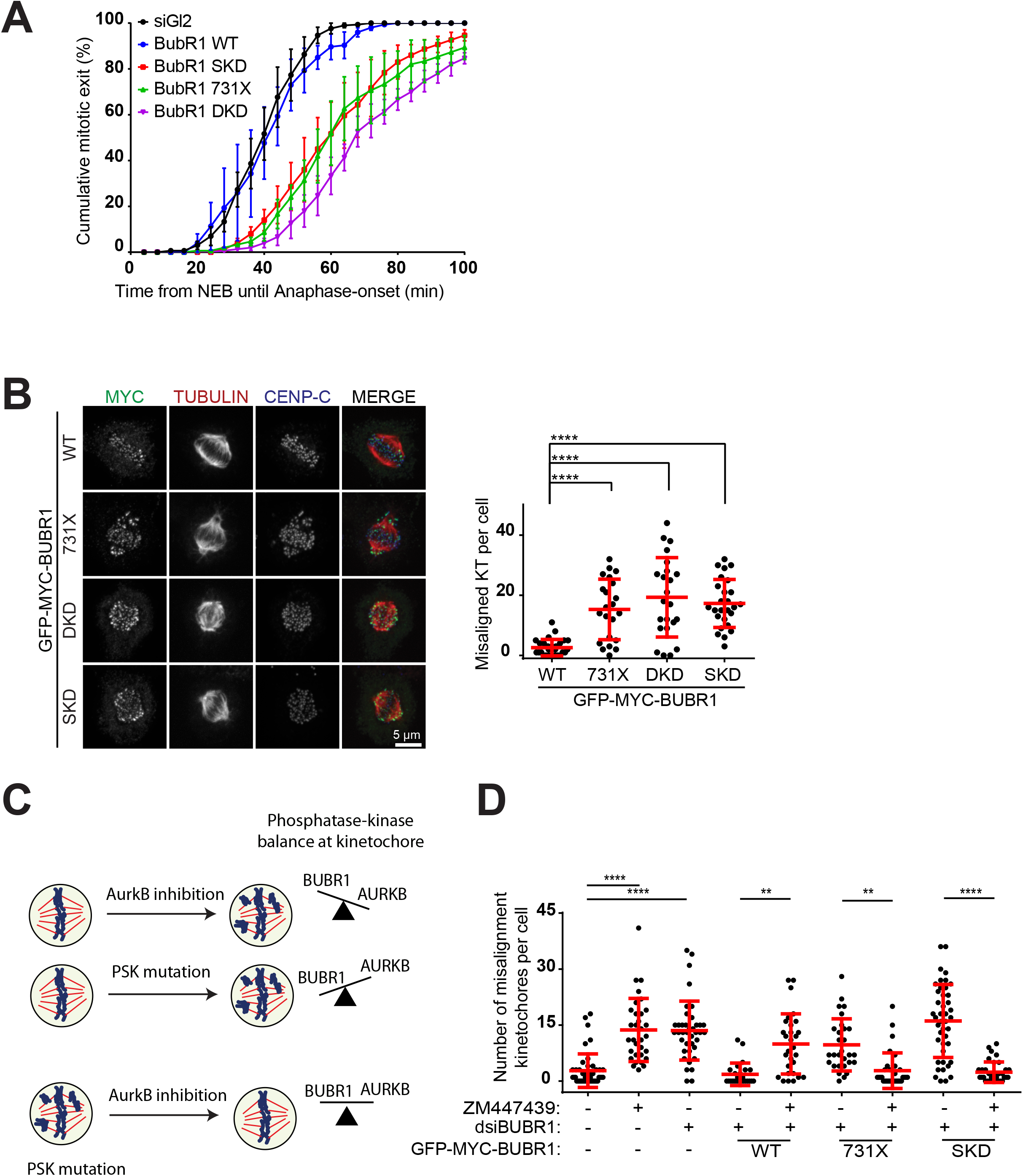
BUBR1 pseudokinase mutants undermine PP2A-B56 functions during mitosis. A: The cumulative mitotic exit from nuclear envelope breakdown (NEB) until anaphase in cells expressing BUBR1 pseudokinase domain mutations at similar levels. Cells were synchronized with thymidine for 24h, and 9h after release, cells were treated with nocodazole and reversine before imaging (n=3, ≥50 cells per experiment). B: Immunofluorescence and quantification of chromosome alignment in cells expressing distinct BUBR1 pseudokinase mutations arrested at prometaphase with STLC for 15h followed by release into MG132 for 2.5h. Fixed cells were stained with anti-CENP-C, anti-tubulin, and anti-MYC. C: Aurora B and BUBR1-PP2A-B56 determine the kinase-phosphatase equilibrium and chromosome alignment at kinetochores in metaphase. Aurora B inhibition favours kinetochore dephosphorylation, which compromises chromosome alignment in cells expressing BUBR1-WT. BUBR1 pseudokinase mutations would disturb phosphatase signalling, increasing Aurora B activity and also undermining alignment. Aurora B inhibition in cells expressing pseudokinase mutants is expected to rescue proper chromosome alignment. D: Graph of chromosome alignment experiment described in (C). Cells expressing BUBR1 WT or pseudokinase mutations and depleted of endogenous BUBR1 were treated as in (B) with or without the Aurora B inhibitor ZM447439.

Another important mitotic function of BUBR1-bound PP2A-B56 is to stabilize kinetochore-fibers, and defects in this induce chromosome bi-orientation and end-on conversion errors^23, 24, 28, 60^. However, evidence regarding a role for the BUBR1 pseudokinase in chromosome alignment is inconsistent ^42, 49, 52, 54, 61, 62^, and we would argue that this is likely due at least in part to the varying levels of mutant (in)stability. In an attempt to address this, we measured alignment of kinetochores in MG132 arrested cells after release from prometaphase arrest in our inducible cell lines and asked whether pseudokinase mutants induce alignment errors when expressed at similar levels to BUBR1-WT. We found that BUBR1 pseudokinase mutants, and BUBR1-731X exhibited a consistent increase in the number of misaligned chromosomes (Fig 4B). In agreement with this, we found fewer stable K-fibers, a decrease in the number of kinetochores attached to microtubules and shorter inter-kinetochore distances in BUBR1-731X, -SKD and -DKD compared to BUBR1-WT (**Suppl. Fig 4A-C)**.

BUBR1-PP2A-B56 and Aurora B kinase activity inversely regulate outer-kinetochore phosphorylation to promote productive attachments ^32, 35, 60^. Whereas Aurora B inhibition causes misalignment as a result of too little kinase/too much phosphatase activity in the presence of BUBR1-WT, the loss of PP2A-B56 activity in pseudokinase mutants results in misalignment as a consequence of too much kinase/too little phosphatase (Fig 4B and ^23, 24, 28, 60^). We therefore reasoned that reducing Aurora B activity in pseudokinase mutant expressing cells could restore the kinase-phosphatase equilibrium and decrease the penetrance of chromosome misalignments (Fig 4C). As expected, Aurora B inhibition in metaphase increased the number of misaligned chromosomes in BUBR1-WT expressing cells, but partially rescued the misalignment observed in the pseudokinase mutants and BUBR1-731X (Fig 4D). A similar conclusion was reached when we measured kinetochore levels of Astrin, a marker of stable end-on attachments ^60, 63^ (**Suppl. Fig 4D**).

### The pseudokinase is required *in cis* for KARD phosphorylation

Our data so far suggests that optimal BUBR1 conformational stability imparted by the pseudokinase domain is an important determinant of KARD phosphorylation. We reasoned that substitution of the pseudokinase domain with the kinase domain of its closest paralog hBUB1, to generate a MADBUB protein should have a stabilizing effect, permitting KARD phosphorylation (Fig. 5A) ^38^. However, we find that while MADBUB was expressed at equivalent levels to BUBR1-WT and able to localize to kinetochores, it was unable to rescue KARD S676 phosphorylation (Fig. 5B).

**Figure 5:**
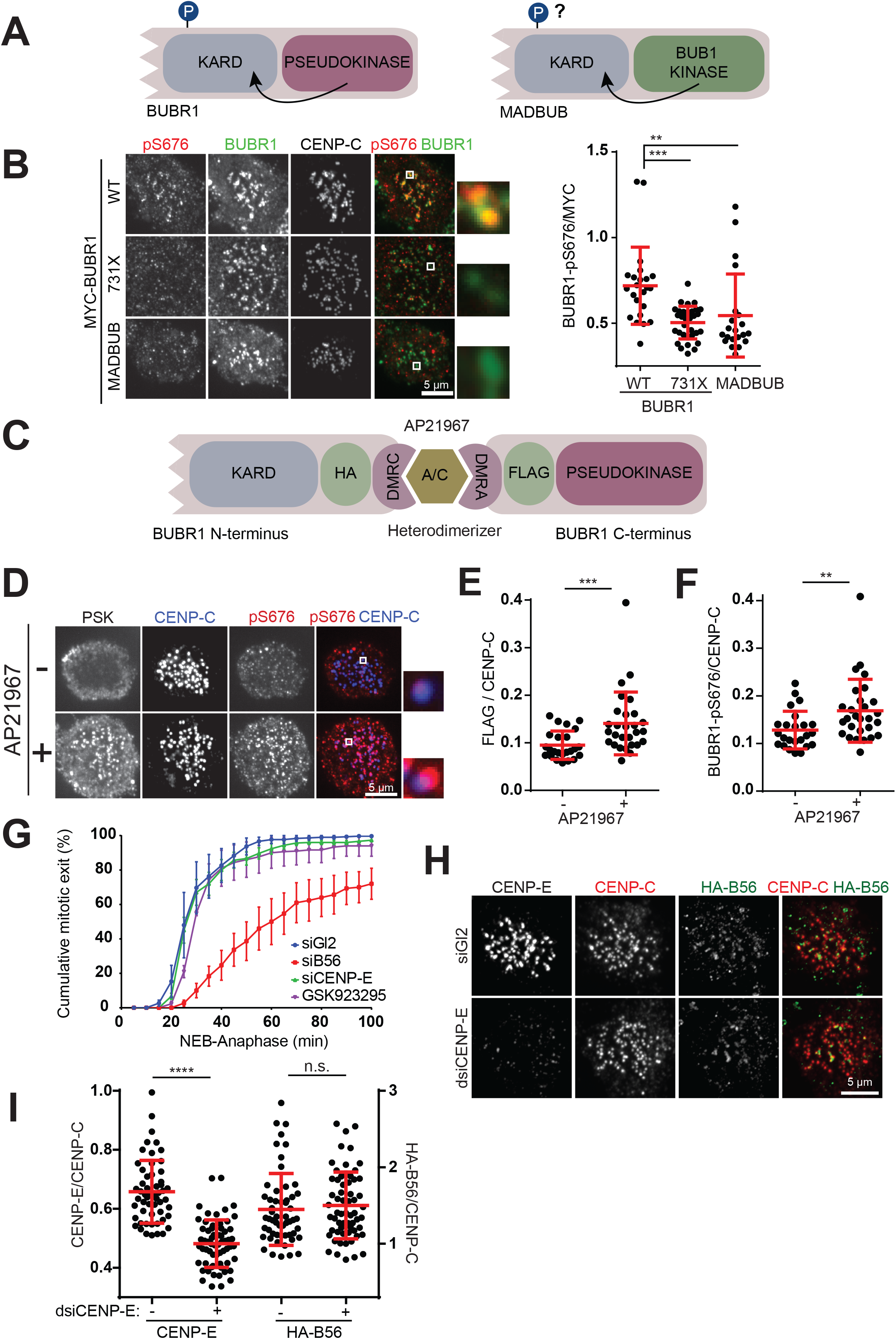
KARD phosphorylation requires the BUBR1 pseudokinase domain to be present in cis. A: A MADBUB-like construct was created by fusing BUBR1-731X to the BUB1 kinase domain (residues 734-1085) to test the requirement for the BUBR1 pseudokinase versus its closest relative BUB1 kinase in KARD phosphorylation. B: HeLa T-Rex cells were transfected with 3xMYC-GFP-BUBR1-WT, -731X, or -MADBUB and arrested in nocodazole for 1 hour before being fixed and stained with the indicated antibodies. The graph shows pS676 levels normalized relative to MYC signals from cells expressing transfected plasmids at similar levels. C: Cartoon of the heterodimerization constructs for N and C-terminal BUBR1. BUBR1 lacking the pseudokinase domain (BUBR1-714X) was fused to DMRA protein at the C-terminus, while BUBR1 (715-1080) was fused to the DMRC at its N-terminus. D: Constructs described in C were transfected in HeLa T-Rex cells depleted of endogenous BUBR1, followed by AP21967 treatment for 5h. During the last hour of treatment, nocodazole was added to the cells. Fixed cells were stained with the indicated antibodies. E-F: Quantification of the BUBR1 C-terminus (E) and pS676 (F) levels at kinetochores from the experiment in D. G: Time-lapse analysis of HeLaS3 GFP-H2B cell line treated with the CENP-E inhibitor GSK923295 (150 nM for 4h) or with GL2, B56 or CENP-E siRNAs. Prior to imaging cells were synchronized with thymidine for 24h and, 9 hours after release, cells were treated with nocodazole and reversine. The graph shows cumulative mitotic index (n=3, ≥50 cells/condition for each experiment). H-I: HeLa T-Rex cells were transfected with HA-B56α and either control GL2 or CENP-E dsiRNA, followed by Nocodazole treatment for 1h. Cells were probed for endogenous CENP-E, CENP-C and HA-B56α. Representative images are shown in H and quantification of kinetochore CENP-E and B56 in I.

To more rigorously test the requirement of the pseudokinase for KARD phosphorylation, we asked whether its presence is required in *cis* or in *trans*, which we tested using drug-induced heterodimerization ^64, 65^ (Fig 5C). When expressed in cells in the absence of the small molecule heterodimerizer AP21967, the C-terminal fragment encoding for the BUBR1 pseudokinase did not localize to kinetochores, consistent with observations that kinetochore tethering of BUBR1 is mediated principally by its N-terminus ^12, 15, 66–70^. Treatment with AP21967 resulted in kinetochore localization of the pseudokinase and concomitant S676 phosphorylation (Fig. 5D-F). We interpret this to mean that the KARD and pseudokinase domains need to be on the same polypeptide (or at least in very close proximity) in order to achieve phosphorylation.

The requirement for the pseudokinase and KARD to be present in *cis* then led us to hypothesize that a pseudokinase-specific interactor may be required for KARD phosphorylation. The only known interacting partner of the BUBR1 kinase domain is the kinesin motor Centromeric protein-E (CENP-E) ^41, 71, 72^. If CENP-E is indeed required for pseudokinase domain stability and KARD phosphorylation, then its depletion or inactivation would be expected to attenuate SAC silencing and diminish B56 recruitment to kinetochores. To test this, we either inhibited its motor activity using GSK923295 or depleted it by siRNA, and observed mitotic exit after MPS1 inhibition by reversine in nocodazole-treated cells as in Fig. 4A. Although we found a significant delay in SAC silencing when B56 isoforms were depleted, neither CENP-E inhibition nor its depletion had any influence on the kinetics of SAC exit, and these cells were indistinguishable from controls (Fig 5G). In full agreement with this, we find that CENP-E depletion did not impact B56 kinetochore recruitment (Fig 5H, I).

The finding that CENP-E played no discernable role in B56 recruitment and SAC silencing was surprising, considering the effect of its inhibition on BUBR1 hyperphosphorylation documented by us and others ^41–43, 73^. However, when we examined KARD phosphorylation in CENP-E depleted cells further, we found that while S676 phosphorylation was reduced, as we have shown before, S670 phosphorylation was increased (^52^ and **Sup Fig 5A-B**). S670 phosphorylation has been shown to be the more important of the two sites for B56 binding, which likely explains why we find no strong requirement for CENP-E in B56 localization and SAC extinction ^24, 27^. Moreover, the interaction between BUBR1 and CENP-E was only appreciably disrupted in BUBR1-731X but not in BUBR1-SKD and -DKD -expressing cells (**Sup Fig 5C**). Therefore, we conclude that pseudokinase-mediated coordination of KARD phosphorylation is not CENP-E dependent.

### BUB1 kinase-BUBR1 pseudokinase heterodimerization drives KARD phosphorylation

Kinase-pseudokinase heterodimerization can exert allosteric regulation on their respective activities ^74–76^. We therefore reasoned that such cooperation may exist between BUB1 and BUBR1. We first tested whether their interaction is required for KARD phosphorylation independently of its function in BUBR1 kinetochore targeting by fusing BUBR1 directly to MIS12, which is recruited constitutively to kinetochores. Indeed, 3xMYC-MIS12-BUBR1 was localized to kinetochores in both control and BUB1-depleted cells, but we consistently observed a reduction in S676 phosphorylation in BUB1-depleted cells compared to controls (Fig 6A; **Suppl. Fig 6A**). Expression of truncated BUB1 lacking the C-terminal kinase domain (BUB1-ΔKIN) also lead to reduced BUBR1 S676 phosphorylation levels (Fig 6B). To confirm that the kinase domain of BUB1 regulates KARD phosphorylation through the pseudokinase domain rather than another region of BUBR1, we tested KARD phosphorylation in cells expressing BUBR1-731X, together with BUB1-WT or BUB1-ΔKIN. Because BUBR1-731X is hyperstable and can be expressed at levels even higher than BUBR1-WT, low-level residual phosphorylation at S676 could be detected when this protein was maximally expressed with doxycyline. Our results demonstrate that S676 phosphorylation of BUBR1-731X remains unchanged between cells expressing BUB1-WT or BUB1-ΔKIN demonstrating that BUB1 regulation of KARD phosphorylation requires the pseudokinase domain (Fig 6C). Finally, given that BUB1 and BUBR1 heterodimerize, we investigated whether an allosteric regulation of BUB1 autophosphorylation by the BUBR1 pseudokinase can also occur. To circumvent the role of the pseudokinase in PP2A-B56 recruitment and thus feedback signalling of BUB1 kinetochore docking ((Fig. 3B, ^31, 35^), we compared the effect of truncating the pseudokinase domain on BUB1 activity in BUBR1 constructs lacking all KARD phosphorylation sites (KARD3A, KARD3A-731X; **Suppl. Fig 6B, C**). To evaluate BUB1 kinase activity, we used phosphospecific antibodies targeting the BUB1 autophosphorylation site T589 ^77^. Loss of the pseudokinase domain attenuated autophosphorylation of BUB1 at T589, in support of the notion that optimal BUB1 phosphorylation requires interaction with the BUBR1 pseudokinase domain (Fig 6D). In accordance with these results, we find that the BUBR1 KARD co-evolved with the BUB1 kinase domain, second only to the BUBR1 pseudokinase domain (**Suppl. Fig. 6D**).

**Figure 6:**
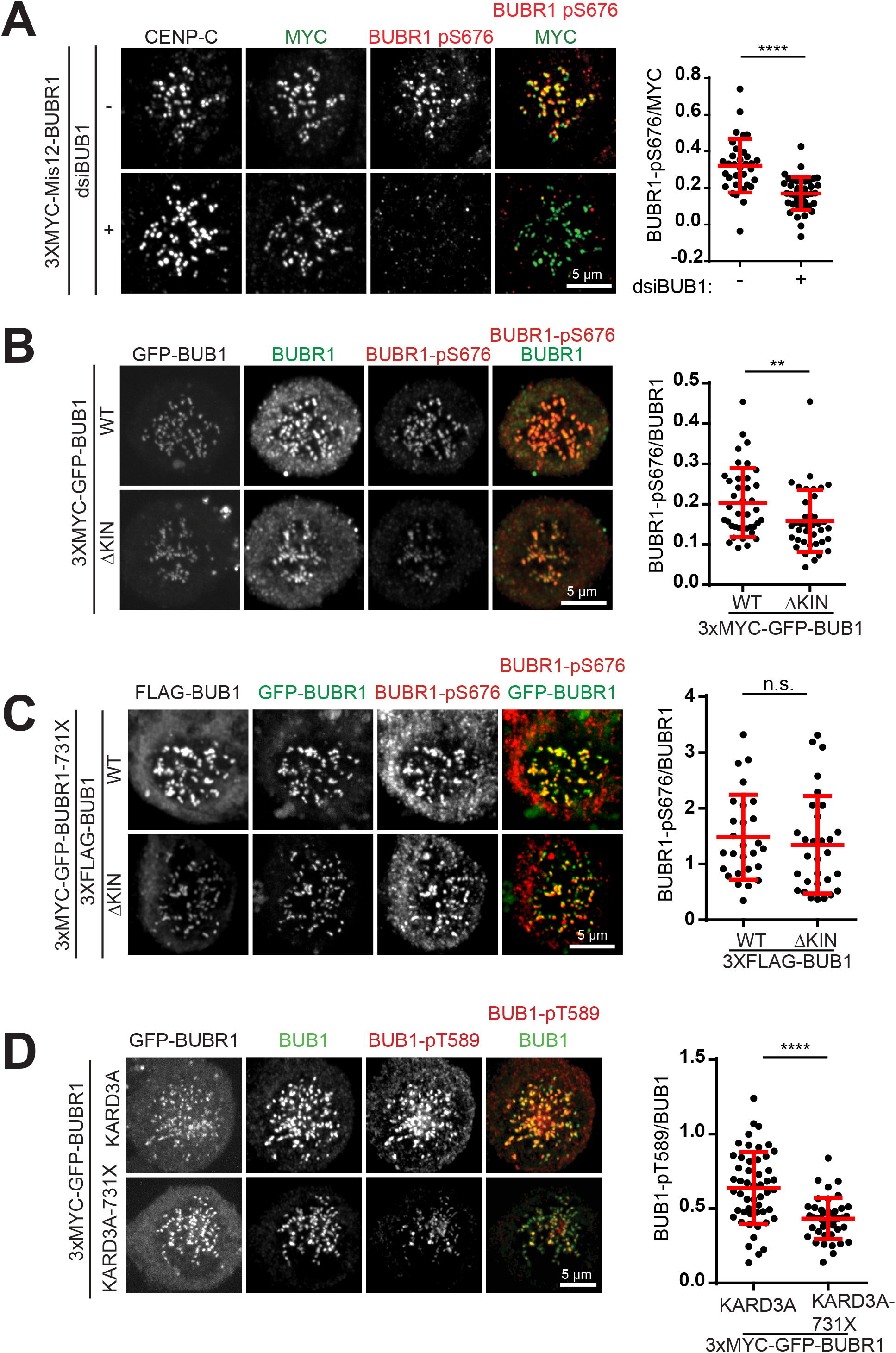
BUBR1 and BUB1 C-termini allosterically regulate each other. A: 3XMYC-Mis12-BUBR1-WT was transfected in Hela-T-REX cells alongside GL2 siRNA or BUB1 siRNAs. Cells we synchronized with thymidine for 8h and released into nocodazole overnight before being fixed and stained with the indicated antibodies. The graph on the right shows relative BUBR1-pS676 levels. B: 3XMYC-BUB1-WT or -ΔKIN were transfected into Hela T-REX cells and together with BUB1 siRNAs. Cells were synchronized with thymidine for 8h and released into nocodazole overnight before being fixed and stained as indicated. The graph on the right shows relative BUBR1-pS676 levels. C: Stable Hela T-Rex cell lines expressing 3xMYC-GFP BUBR1-731X were transfected with 3xFLAG-BUB1-WT or ΔKIN together with BUB1 and BUBR1 dsiRNAs. Cells were synchronized with thymidine for 8h and released into nocodazole overnight before being fixed and stained as indicated. The graph on the right shows relative BUBR1-pS676 levels. D: Stable Hela T-Rex cell lines expressing 3xMYC-GFP BUBR1-KARD3A or KARD3A-731X were transfected with BUBR1 siRNAs, synchronized with thymidine for 8h and released into nocodazole overnight. Cells were subsequently fixed before staining with the indicated antibodies. The quantitation on the right shows relative kinetochore levels of BUB1 pT589.

## Discussion

Whether BUBR1 is a *bona fide* active kinase is a subject that has generated considerable controversy ^38, 41–43, 73, 78^. If BUBR1 is indeed inactive as proposed before and as we confirm here, an important question that has not yet been addressed is whether the BUBR1 pseudokinase domain retained some function of the ancient MADBUB kinase domain, or acquired new activity as a result of neofunctionalization, that might explain its conservation during vertebrate evolution. One hypothesis put forward is that in the case of hBUBR1, the pseudokinase was retained as it provides structural stability to the protein ^38–40^. However, while this is indeed true, stability alone cannot explain the retention of the kinase fold because losing the pseudokinase domain entirely stabilizes BUBR1 protein levels ^49^. Furthermore, that the pseudokinase is conserved in complex eukaryotes raises the possibility that it may have evolved unique functions in this context ^12^. The collective phylogenomic, biochemical and functional evidence presented here, definitively demonstrates that hBUBR1 is indeed a pseudokinase, although we cannot formally exclude catalytic ability under conditions we have not explored.

Strikingly, we demonstrate that the presence of the intact pseudokinase domain in the BUBR1 polypeptide is required for efficient phosphorylation of the KARD and PP2A-B56 recruitment to the outer kinetochore. As a consequence, the pseudokinase domain coordinates critical mitotic events such as silencing of SAC signals and the counterbalance of Aurora B activity at the outer kinetochore. The pseudokinase presumably functions through allosteric interaction with the C-terminal BUB1 kinase domain. Indeed, the BUBR1 pseudokinase and BUB1 kinase domains appear to allosterically fine-tune phosphorylations on each other in their respective C-termini: KARD phosphorylation is reduced in cells expressing BUB1 lacking the kinase domain and BUB1 autophosphorylation at T589 is likewise partially lost in BUBR1-731X expressing cells. Therefore, in humans, maximal KARD phosphorylation is obtained when both BUB1 kinase and BUBR1 pseudokinase domains are present. This also suggests that in addition to the major BUB1-BUBR1 dimerization interface in their respective N-termini, interactions could also occur at the kinase-pseudokinase level. In support of this idea BUB1 truncated at the C-terminus failed to fully maintain BUBR1 at kinetochores and the major BUBR1 binding region in BUB1 (residues 209-409) exhibited a modest reduction in co-immunoprecipitated BUBR1 compared to full-length BUB1 ^12^. A three-dimensional structure of the BUB1-BUBR1 heterodimer will undoubtedly be a critical step towards understanding the molecular details of this interaction.

Notably, this regulation of BUB1 activity by the BUBR1 pseudokinase may also explain the kinase activity reported in BUBR1 immunoprecipitates, and why this activity is sensitive to mutation of BUBR1 catalytic triad residues, or loss of the BUBR1 kinase which we have shown here tempers BUB1 autophosphorylation ^41–43, 73^. In agreement with this, BUB1 and BUBR1 form relatively stable complexes *in vitro* ^44^, and previous work has shown that BUBR1-associated kinase activity is particularly sensitive to BUB1 inhibitors ^38^. Importantly, the BUBR1 pseudokinase domain functions as a modulator rather than an ON-OFF switch for BUB1 kinase activity, as would be predicted by its conservation in only a number of species. Previous work from the Musacchio group suggested that in vitro, full-length BUBR1 does not impinge upon H2A phosphorylation by BUB1 ^44^. Indeed, we did not observe a consistent reduction of H2A phosphorylation when the BUBR1 pseudokinase was mutated or removed (data not shown). H2A-T120 is a particularly strong substrate for BUB1 and small changes in phosphorylation may not be readily detectable by the methods we used here. Alternatively, it is plausible that only changes on the BUB1 polypeptide itself are appreciably regulated by the heterodimer interaction.

Kinase-kinase and kinase-pseudokinase allosteric regulation, where the role of the pseudokinase is to assist the bona fide kinase in the pair into an active conformation, is not unprecedented. Indeed, it has been argued that the conformational plasticity of the kinase fold has served two purposes throughout evolution and as such, in all kinases: the phosphotransfer reaction and noncatalytic scaffolding or allosteric functions ^75, 79–81^. Without structural information we can only speculate on the mechanism of how BUBR1 kinase conformation is conduit to KARD phosphorylation and allosteric regulation of BUB1. Because BUBR1 requires the presence of canonical catalytic residues for stability, its adoption of an active-like kinase conformation may either relieve steric constraints or drive allosteric conformational changes in the KARD and BUB1 C-terminus allowing for their optimal phosphorylation and activation. This type of mechanism has been previously identified in other pseudokinases that function as activators of the partner kinase including JAK2, STRADα and HER3 ^82–85^.

Finally, our work here may also shed light on the evolutionary history of BUBR1. Previous phylogenomic studies indicated that MAD3/BUBR1 orthologs evolved through numerous gene duplication and subfunctionalization events ^38–40^. Our data support this idea and further invoke the hypothesis that the ancient MADBUB possessed both protein kinase activity and a non-catalytic allosteric function, and may indeed have functioned as a dimer, very much like many modern day kinases. In this scenario, following the MADBUB gene duplication event with the emergence of vertebrates, one gene (BUBR1) relinquished phosphoryl-transfer function but retained some functionality as an allosteric activator of its kinase-active partner (BUB1). Because the KARD is only present in a subset of BUBR1 orthologs, the pseudokinase domain may have been co-opted to stabilize the KARD-PP2AB56 interaction more recently during evolution. The evolutionary pressure to fine-tune SAC signalling and coordinate PP2A and PP1 activities more precisely in complex eukaryotes may have eventually evolved to be the more critical function of the pseudokinase ^2^.

## Material and Methods

### Evolutionary analysis

#### Sequence retrieval

BUB1 (BUB) and BUBR1 (MAD) sequences were taken from the dataset compiled by Tromer et al ^39^ and from Ensembl Orthologs ^50^ based on a search of orthologs for the human BUB1 and BUBR1 sequences, and downloaded on December 5th, 2018. Sequences from the different data sets that were annotated to the same organism and shared over 95% sequence identity when aligned using BLASTP 2.6.0+ ^86^ were considered to be redundant, in which case the Ensembl annotated sequences were retained. The complete data set includes 173 sequences for BUB1 and 176 for BUBR1. All sequences are available at https://github.com/landrylaboratory/BuBR1_coevolution.

#### Domain annotation

Domain annotations for BUB1 and BUBR1 sequences were taken from Tromer et al. Using these sequences as a reference, the corresponding domains were annotated in the sequences from Ensembl Orthologs using jackhmmer (HMMER suite version 3.1b2) ^87^. Hits with a posterior probability of >0.95 and a coverage of >60% of the length of the reference domains were maintained. Hits were considered to overlap if the overlap represented >50% of the total bases or >70% of the shorter sequence. Following Geer et al. ^88^, the best-scoring hits were maintained as the final domain annotation. Copresence of protein domains was analyzed by Pearson correlation as in ^39^ after discretizing the annotations to reflect absence (0) or presence (1) of a specific domain in each sequence.

#### Coevolutionary analysis

Multiple sequence alignments were performed with the Multiple Domain Alignment Tool (MDAT) version 1.0 ^89^ using the final annotation derived above (Supplementary Files). Using the Evol library from the ProDy package version 1.10.8 ^90^, the alignments were then refined using the human sequences as the reference and filtered for 80% occupancy, that is, only sequences for which gaps represented 20% or less of the full sequence were retained. Finally, only organisms for which both BUB1 and BUBR1 had passed the filtering process were included. This yielded a dataset of 77 sequences for each of BUB1 and BUBR1.

Shannon entropy as a measure of per site diversity and mutual information between pairs of residues were calculated with the Evol suite ^90^ following the definitions provided by Martin, et al ^91^. Mutual information was then corrected for background noise using the average product correction defined by Dunn et al. ^51^. Coevolution between pairs of residues was extended to the domain level by calculating the mean of the MIp of all possible pairs of residues for a given pair of domains and normalizing by the MIp of a domain with itself, since mutual information is dependent on the entropy of each pair of domains. This analysis was carried out with BUB1 and BUBR1 separately as well as with their concatenated sequences. Domains that were not present in the human paralogs (D-Box, MadaM for BUB1; CMI, MadaM, and “Other KEN” for BUBR1) were excluded from the analysis. The “Other KEN” domain for BUB1 was also excluded because it could not be confidently retrieved by jackhmmer and it overlapped with the “Other ABBA” domains. Reference positions of the domains for the human sequence were taken from Tromer, et al. ^39^, accounting for a region of 14 residues within the TPR domain of the human BUBR1 sequence in that data set that was missing in the sequence from Ensembl orthologs.

All scripts used for the domain annotation and the analysis of domain co-evolution are available at: https://github.com/landrylaboratory/BuBR1_coevolution

#### Cell culture and transfection

Cell lines were cultured in DMEM (Hyclone) containing 10% (vol/vol) of fetal bovine serum and Pen/Strep (100 μg/ml, Hyclone) at 37°C and 5% CO_2_. Plasmids and dsiRNAs were transfected using JetPrime (Polyplus), Interferin (Polyplus) or Oligofectamine (Invitrogen) according to the manufacturer’s instructions. Stable HeLa cell lines were generated using the T-Rex doxycycline-inducible Flp-In system (Life Technologies), selected with Hygromycin B (Wisent, Cat# 450-141-XL, 2 mg/mL), and induced with Tetracycline (Sigma, 24h concentration spanning from 25 ng/mL to 1 μg/ml). Drug treatments were performed as follows unless otherwise indicated: thymidine (Acros Organics, 2mM for 8 h), MG132 (Calbiochem, 20 µM for 2.5 h), STLC (Sigma, 5 µM, 12h), ZM-447439 (Enzo, 2 μM, 30 mins), GSK923295 (Selleckchem, S7090, 150 nM), Nocodazole (Sigma, Cat# M1404-10MG, 100 ng/mL for 12 h), A/C Heterodimerizer AP21967 (Takara Bio, 500 nM for 5 hours), Okadaic Acid (Santa Cruz Biotechnology, SC-202259A, 1µg/mL, 1 hour), and Reversine (Sigma, Cat# R3904-1MG, 1 mM for 30 minutes). Knock-downs were obtained with dsiRNA (IDT) and analyzed 24-78h after transfection. The dsiRNAs target the following sequences in the target proteins as indicated: dsiCENP-E (used at 50nM) 5’-CUCUUACUGCUCUCCAGUUUGCCAG -3’; dsiBUB1 (used at 20 nM each to target siRNA-resistant constructs) 5’-GAGUGAUCACGAUUUCUAAAUCAGA-3’ and 5’-CCAGUGAGUUCCUAUCCAAAUACUU-3’; dsiBUBR1 (20-50 nM) sequence is 5’-GUCUCACAGAUUGCUGCCUCAGAGC -3’; siB56 target sequences were previously reported ^32^ and were mixed in an equimolar concentration to generate a stock solution of 18.2 uM for each subunit except for B56ε, which was at 27.3 uM. dsiB56 treatments were performed at a final concentration of 45 nM. B56 target sequences are as follows; B56α: 5’-GAAACAUACUCAACCAGUUCAUUCA-3’, B56β: 5’-GAGGCUUAGGAUAUACUCAUUGUUC-3’, B56γ: 5’-UGCUUCUAACGUUGGUUCAUCUUCC-3’, B56δ: 5’-AGUCCCACAAUUACCGGCUCAGUCA-3’, and B56ε: 5’-GUACUAUACAAUAUGCCAGCUGUGC-3’.

#### Plasmid construction

All point mutants described here were generated through site-directed mutagenesis and were verified by sequencing. For the stable cell lines, 3XMYC-GFP-BUBR1-WT was inserted into BamH1-Not1 sites in pcDNA5/TO/FRT from pcDNA 3.1-3X-MYC-BUBR1-WT ^53^. The HA-B56α plasmid (pCEP-4HA-PP2A-B56α) was purchased from Addgene (ref #14532). All BUB1 constructs were cloned into pcDNA5/TO/FRT vectors at BamH1-Not1 sites. The MADBUB plasmid was generated using Gibson cloning using the NEBuilder® HiFi DNA Assembly Master Mix (New England BioLabs) following the manufacturer’s instructions. Essentially, residues 734-1085 from BUB1 were cloned C-terminal to the BUBR1 sequence in pcDNA5/TO/FRT 3XMYC-GFP-BUBR1-731X, while the stop codon in BUBR1-731X was silenced. Vectors for the DMRC/DMRA system (pcDNA5/FRT/TO-HA-DMRC and pcDNA5/FRT/TO-FLAG-DMRA) were a gift from Amélie Fradet-Turcotte. BUBR1 residues 1-714 was inserted at cloning sites AflII-KpnI (pcDNA5/FRT/TO-BUBR1-714X-HA-DMRC), while BUBR1 residues 715-1050 were inserted at the NotI-ApI sites (pcDNA5/FRT/TO-FLAG-DMRA-Pseudokinase).

#### Antibodies and dye

The following antibodies were used in this study at 1mg/mL, unless otherwise stated: anti-CENP-C (PD030, MBL International), mouse anti-CENP-E (1H12, Abcam), rabbit anti-CENP-E (C7488, Sigma), anti-H2A-pT120 (61195, Active Motif), anti-α-TUBULIN (DM1A, Santa Cruz), anti-MYC (9E10, Thermo Scientific), anti-HA (CloneHA7, H9658, Sigma), anti-FLAG (M2, F1804, Sigma), CREST anti-centromere serum (HCT-0100, Immunovision), and anti-ASTRIN (NB100-74638, Novus). The antibodies rabbit BUBR1-pS676 ^52^, mouse monoclonal BUBR1 ^52^, rabbit BUBR1-pS670 ^53^, anti-BUB1 ^53^, and rabbit BUB1-pT589 ^77^ were previously described. Anti-KNL1 pS24 was a kind gift from Iain Cheeseman.

For immunofluorescence, AlexaFluor-AffiniPure series secondary antibodies (Jackson ImmunoResearch) were used at 1:1000 and, for Western blotting, horseradish peroxidase-coupled secondary antibodies (Jackson ImmunoResearch) at 1:10000. For live-cell imaging, cells were washed with PBS and treated with SiR-DNA dye (CY-SC007, Cytoskeleton) at 300 nM for 6 hours before starting imaging.

#### Immunofluorescence

Cells were seeded on cover slips and subsequently treated with the appropriate dsiRNA after adhesion, followed by synchronization and drug treatment as indicated. Cells were either simultaneously fixed and extracted by PTEMF (0.2% Triton X-100, 20mM PIPES pH 6.9, 1mM MgCl_2_, 10mM EGTA and 4% formaldehyde) or by pre-extraction (1% Triton X-100, 25mM PIPES pH 6.9, 2mM MgCl_2_, 10mM EGTA) followed by fixation (25mM PIPES pH 6.9, 2mM MgCl_2_, 10mM EGTA and 4% formaldehyde), depending on the antibody. After fixation, cells were blocked in 3% BSA in PBS and incubated with antibodies for 1h at room temperature or 24h at 4°C, and then secondary antibodies for 1-2h at room temperature.

#### Immunoprecipitation and Immunoblotting

The mitotic fraction of cells was collected by shake-off, washed with PBS and lysed with RIPA lysis buffer (Tris-HCL 150 mM pH 7.5, NaCl 150 mM, NaF 10 mM, NP-40 1%, Na-deoxycholate 0.1%, B-glycerophosphate 20 mM, sodium vanadate 0.1mM, sodium pyrophosphate 10mM, Leupeptin 1 mg/mL, 1 mg/mL Aprotinin, and 1mM AEBSF). Cells were incubated with lysis buffer for 30-45 minutes under agitation at 4°C, followed by 14 000 g centrifugation at 4°C for 10 minutes to clear the lysates. The soluble fraction was collected, and protein content was measured by BCA assay (Thermo Scientific). For immunoprecipitation, cell lysates were incubated overnight with 10 µL Protein G Agarose beads at 4°C and anti-MYC antibody. The beads were then washed with lysis buffer at least 3X, boiled with Laemmli buffer and resolved by SDS-PAGE. Immunoblotting was performed according to standard protocol and imaged with a Bio-Rad ChemiDoc MP.

#### Microscopy

Immunofluorescence and live-cell images were acquired with an Olympus IX80 on an inverted confocal microscope equipped with a WaveFX-Borealin-SC Yokagawa spinning disc (Quorum Technologies) and an Orca Flash4.0 camera (Hamamatsu). The system was equipped with a motorized stage (ASI) and incubator with atmospheric CO_2_ warmed at 37°C for live-cell imaging. We used Metamorph software (Molecular Devices) to acquire images and ImageJ2 (Fiji version 1.52i) to analyze and process the data. For time-lapse experiments, images were taken with a 20X objective every 4-5 minutes, and only cells expressing GFP-BUBR1 were included in the analysis.

#### Image analysis

The kinetochore signal intensity was measured according to protocol in published literature ^92^ in ImageJ2 (Fiji version 1.52i). Briefly, cells were measured individually by cropping the desired cell, creating a selection of the kinetochores area by thresholding, and transferring to and measuring the selection in the desired channels. Whole spindle intensity measurements after cold-induced depolymerization and inter-kinetochore distance analysis were performed according to a published protocol ^93^, where tubulin intensity at the spindle was subtracted from the tubulin average of 6 distinct 10×10 pixel square measured inside the cell but outside the spindle. KT-MT attachment levels were measured also by selecting the kinetochore area and transferring the selected region to the tubulin image. The number of misaligned kinetochores was measured by equally dividing the area in between the poles of the cell in 5 zones. The middle zone was considered the alignment zone and was excluded from further analysis. The remaining misaligned kinetochores were then selected by the Huang auto-threshold function in ImageJ2. The image was then processed using functions “Make binary”, “Watershed”, and “Analyze Particles”.

#### Statistical Analysis

Unless otherwise stated, all experiments and statistical analysis were performed on triplicate experiments. Statistical analysis and graphic plotting were performed in GraphPad Prism Software V6.01. If the experiment comprises only two conditions, non-parametric t-test (Mann Whitney test) was used to determine difference significance. In experiments comprising more than two conditions, test significance was calculated by non-parametrical ANOVA (Kruskal-Wallis test), and conditions compared using Dunn’s multiple comparisons test. *P*-values smaller than 0.005 are considered significant, where * for *P* ≤ 0.05, ** for *P* ≤ 0.01, *** for *P* ≤ 0.001, **** for *P* ≤ 0.0001.

## Acknowledgements

We would like to thank members of the Elowe lab for support and discussion, Dr. Isabelle Lucet for helpful discussions, and Drs. Amélie Fradet-Turcotte and Iain Cheeseman for gifting reagents. We would also like to thank Joanie Patenaude for technical assistance. This work was supported by a CIHR project grant to SE (RN376557) and CIHR Foundation Grant to CRL (RN348479 - 387697). SE hold an FRQS J2 salary award and CRL the Canada Research Chair in Evolutionary Cell and Systems Biology. LGB has been supported by training awards from Desjardins, PROTEO, and “Centre de Recherche sur le Cancer de l’Université Laval”.

