## Supplemental material for "The BUBR1 pseudokinase domain promotes efficient kinetochore PP2A-B56 recruitment to regulate spindle checkpoint silencing and chromosome alignment"

Luciano Gama Braga *et al.*

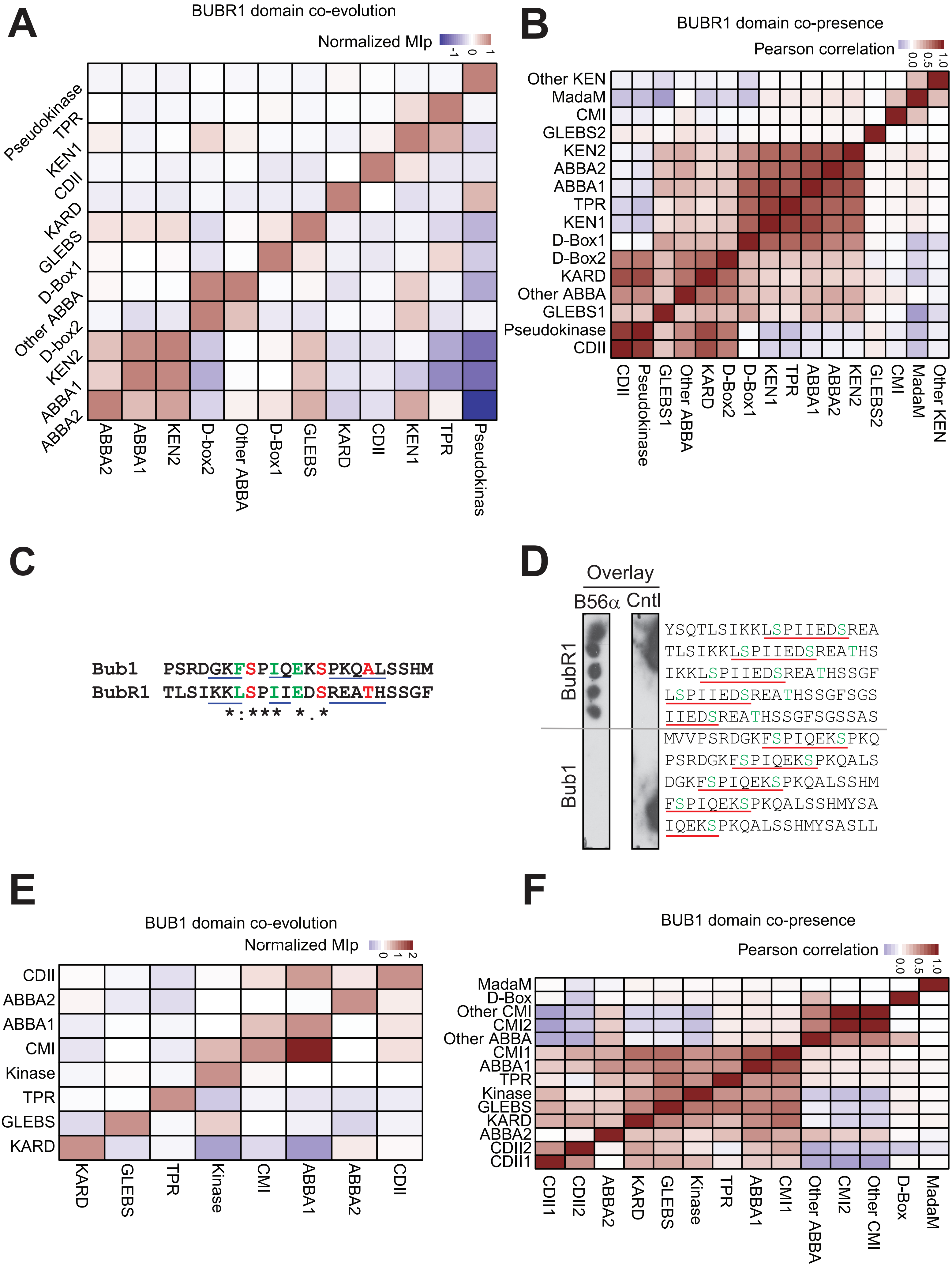
A
BUBR1 domain co-evolution
Normalized MIp
-1
0
1
Pseudokinase
TPR
KEN1
CDII
KARD
GLEBS
D-Box1
Other ABBA
D-box2
KEN2
ABBA1
ABBA2
ABBA2
ABBA1
KEN2
D-box2
Other ABBA
D-Box1
GLEBS
KARD
CDII
KEN1
TPR
Pseudokinas
B
BUBR1 domain co-presence
Pearson correlation
0.0
0.5
1.0
Other KEN
MadaM
CMI
GLEBS2
KEN2
ABBA2
ABBA1
TPR
KEN1
D-Box1
D-Box2
KARD
Other ABBA
GLEBS1
Pseudokinase
CDII
CDII
Pseudokinase
GLEBS1
Other ABBA
KARD
D-Box2
D-Box1
KEN1
TPR
ABBA1
ABBA2
KEN2
GLEBS2
CMI
MadaM
Other KEN
C
Bub1 PSRDGKFSPIQEKSPKQALSSHM
BubR1 TLSIKKLSPIIEDSREATHSSGF
*:*** *.*
D
Overlay
B56α
Cntl
BubR1
Bub1
YSQTLSIKKLSPIIEDSREA
TLSIKKLSPIIEDSREATHS
IKKLSPIIEDSREATHSSGF
LSPIIEDSREATHSSGFSGS
IIEDSREATHSSGFSGSSAS
MVVPSRDGKFSPIQEKSPKQ
PSRDGKFSPIQEKSPKQALS
DGKFSPIQEKSPKQALSSHM
FSPIQEKSPKQALSSHMYSA
IQEKSPKQALSSHMYSASLL
E
BUB1 domain co-evolution
Normalized MIp
0
1
2
CDII
ABBA2
ABBA1
CMI
Kinase
TPR
GLEBS
KARD
KARD
GLEBS
TPR
Kinase
CMI
ABBA1
ABBA2
CDII
F
BUB1 domain co-presence
Pearson correlation
0.0
0.5
1.0
MadaM
D-Box
Other CMI
CMI2
Other ABBA
CMI1
ABBA1
TPR
Kinase
GLEBS
KARD
ABBA2
CDII2
CDII1
CDII1
CDII2
ABBA2
KARD
GLEBS
Kinase
TPR
ABBA1
CMI1
Other ABBA
CMI2
Other CMI
D-Box
MadaM

**Suppl Fig. 1**

**Suppl. Fig. 1: Conservation between BUB1 and BUBR1 motifs across evolution**

A: Co-evolution heatmap of individual BUBR1 domains. The normalized mutual information of a protein (MIp) directly correlates to co-evolving positions in protein families.

B: Heatmap of the Pearson correlation between BUBR1 domains as an indicator of domain co-presence.

C: Alignment of the KARD and KARD-like domain of hBUBR1 and hBUB1, respectively.

D: Far-Western of the B56-α binding to the KARD sequences from BUBR1 and BUB1 synthesized as a peptide array. Core B56 binding sequences are underlined in red and residues in green were incorporated as phosphoresidues. Peptide membranes were incubated with GST-B56α, washed and immunoblotted with anti-GST antibodies.

E: Co-evolution heatmap of BUB1 domains. The normalized mutual information of a protein (MIp) directly correlates to co-evolving positions in protein families.

F: Heatmap of the Pearson correlation of BUB1 domains as an indicator of domain co-presence. In parenthesis is the number of sequences where the domain was recognized.

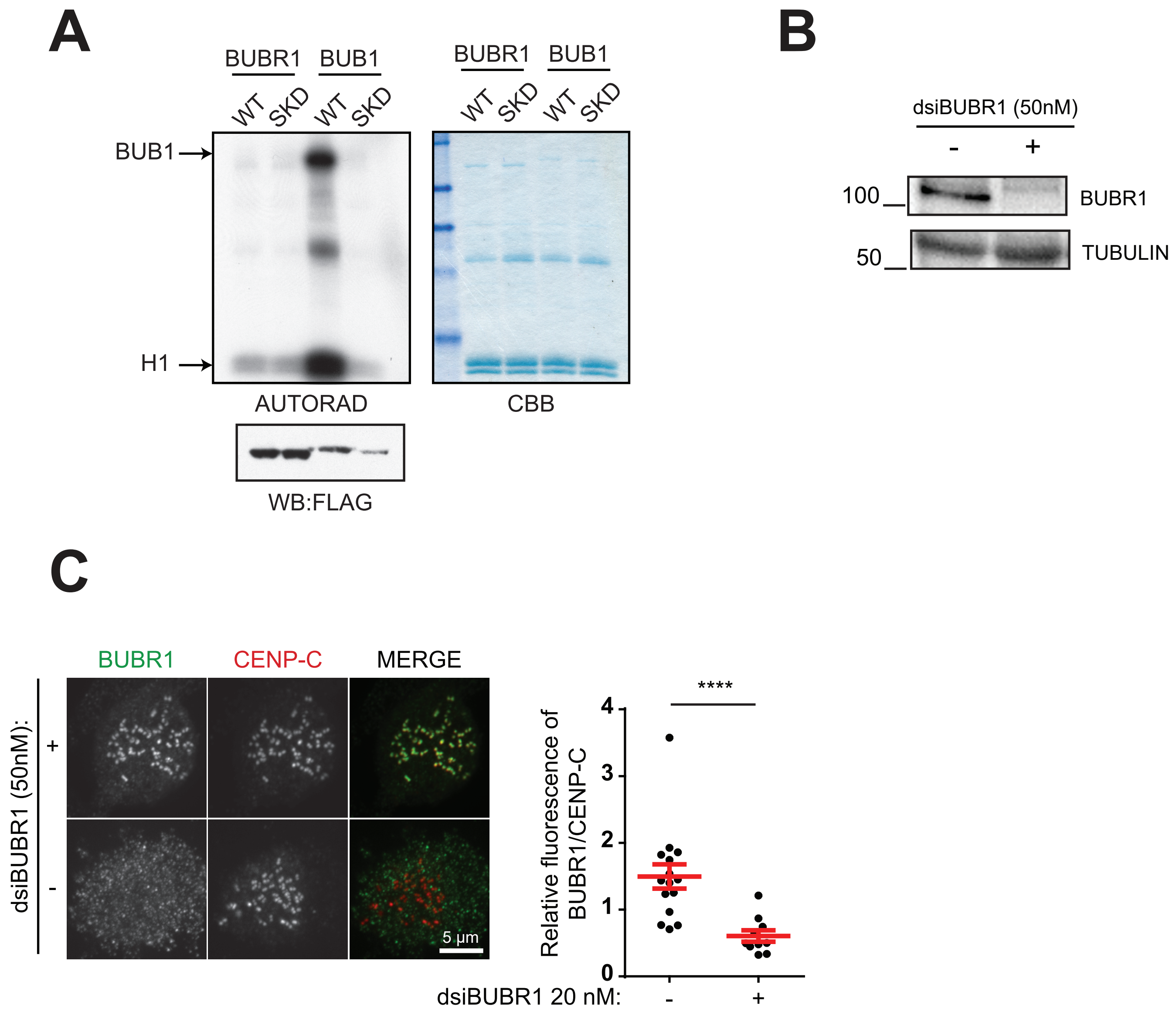
A
BUBR1
BUB1
WT
SKD
WT
SKD
BUB1
H1
AUTORAD
WB:FLAG
BUBR1
BUB1
WT
SKD
WT
SKD
CBB
B
dsiBUBR1 (50nM)
-
+
100
BUBR1
50
TUBULIN
C
BUBR1
CENP-C
MERGE
dsiBUBR1 (50nM):
+
-
5 µm
****
Relative fluorescence of BUBR1/CENP-C
4
3
2
1
0
dsiBUBR1 20 nM:
-
+

**Suppl Fig. 2**

**Suppl. Fig. 2: BUBR1 is a pseudokinase domain that lacks phosphotransfer activity.**

A: In vitro kinase assay of WT and catalytic triad mutants of 3xMYC-tagged BUB1 and BUBR1. BUBR1-WT and SKD as well as BUB1 WT and KD (D946A) were expressed in HEK-293T cells and synchronized in mitosis with nocodazole. 3xMYC-tagged proteins were immunoprecipitated from cleared lysates and subjected to an in vitro kinase assay to monitor autophosphorylation and phosphorylation of the exogenous substrate histone H1.

B-C: Depletion efficiency of BUBR1 in Hela T-Rex cells demonstrated by Western blotting (B) and immunofluorescence (C).

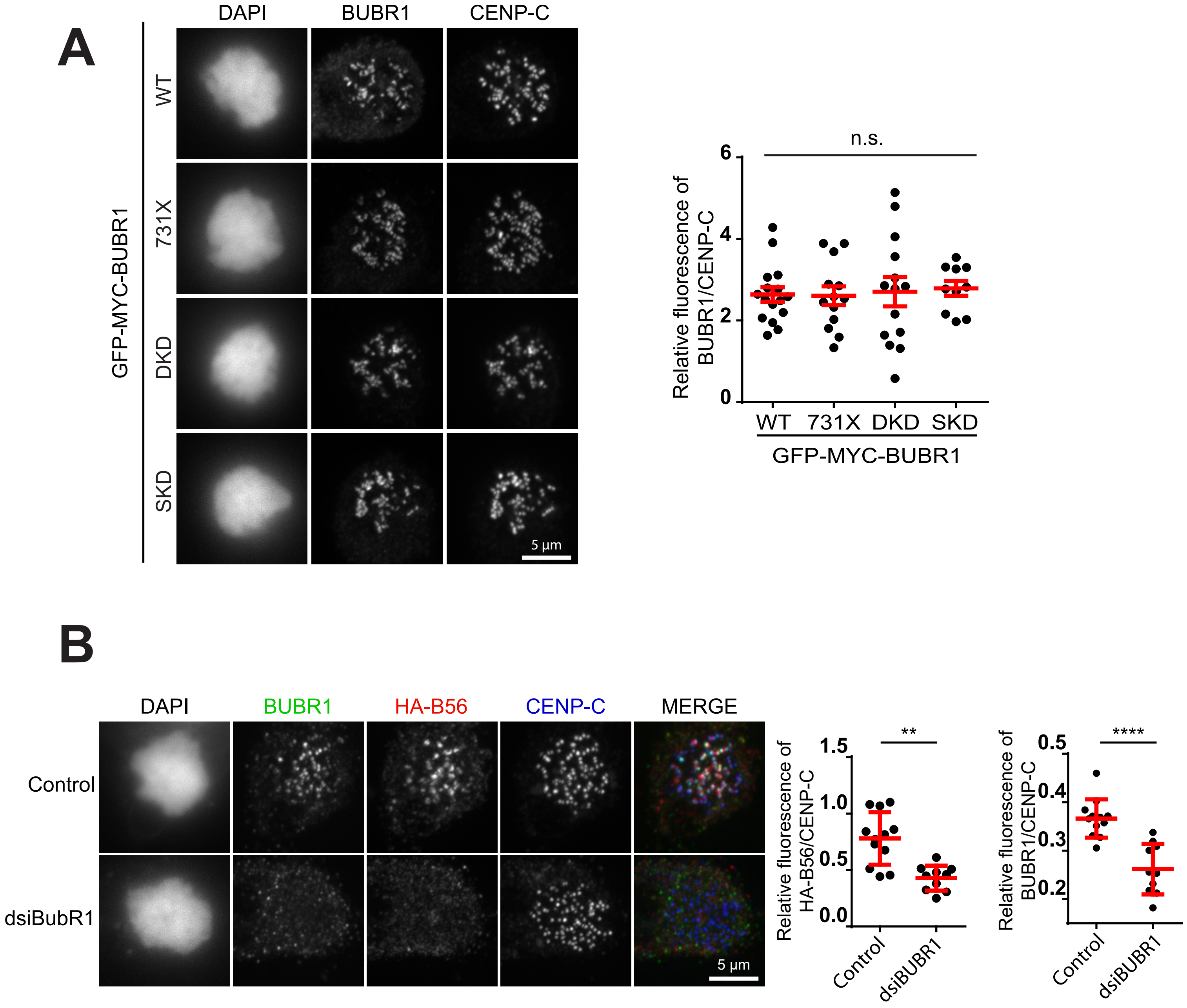
A
DAPI
BUBR1
CENP-C
GFP-MYC-BUBR1
WT
731X
DKD
SKD
5 µm
n.s.
Relative fluorescence of BUBR1/CENP-C
6
4
2
0
WT
731X
DKD
SKD
GFP-MYC-BUBR1
B
DAPI
BUBR1
HA-B56
CENP-C
MERGE
Control
dsiBubR1
5 µm
**
Relative fluorescence of HA-B56/CENP-C
1.5
1.0
0.5
0.0
Control
dsiBUBR1
****
Relative fluorescence of BUBR1/CENP-C
0.5
0.4
0.3
0.2
Control
dsiBUBR1

**Suppl Fig. 3**

**Suppl. Fig. 3: BUBR1 and B56 expression in the Hela T-Rex stable cell lines.**

A: Equivalent expression of BUBR1 mutants in the different stable cell lines used in this study. BUBR1 expression in the cell lines was induced with tetracycline as follows: at 25 ng/mL for BUBR1-WT, 50 ng/mL for BUBR1-731X, 1000 ng/mL for BUBR1-SKD, and 1000 ng/mL for BUBR1-DKD.

B: HeLa T-Rex cells expressing HA-B56α were treated with siRNAs against BUBR1 or GL2 as a control. Cells were treated with nocodazole then fixed and immunostained for BUBR1, HA and CENP-C. Quantitation of relative HA-B56α and BUBR1 kinetochore intensity is shown on the graph on the right.

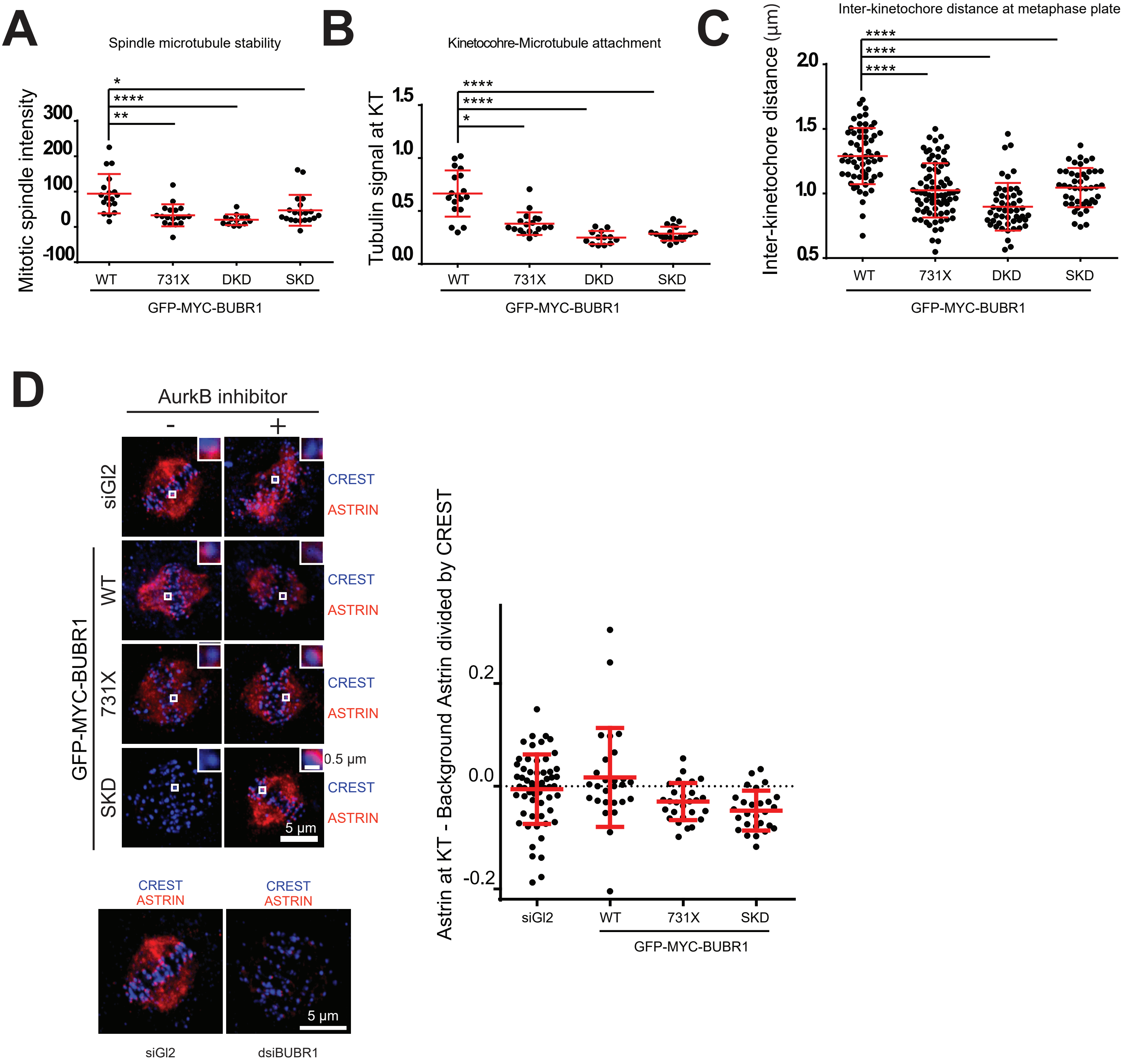
A
Spindle microtubule stability
Mitotic spindle intensity
300
200
100
0
-100
*
****
**
WT
731X
DKD
SKD
GFP-MYC-BUBR1
B
Kinetocohre-Microtubule attachment
Tubulin signal at KT
1.5
1.0
0.5
0.0
****
****
*
WT
731X
DKD
SKD
GFP-MYC-BUBR1
C
Inter-kinetochore distance at metaphase plate
Inter-kinetochore distance (μm)
2.0
1.5
1.0
0.5
****
****
****
WT
731X
DKD
SKD
GFP-MYC-BUBR1
D
AurkB inhibitor
-
+
siGl2
GFP-MYC-BUBR1
WT
731X
SKD
CREST
ASTRIN
0.5 μm
5 μm
Astrin at KT - Background Astrin divided by CREST
0.2
0.0
-0.2
siGl2
WT
731X
SKD
GFP-MYC-BUBR1
CREST
ASTRIN
siGl2
dsiBUBR1
5 μm

Suppl Fig. 4

**Suppl. Fig. 4: Mitotic phenotypes associated with BUBR1 pseudokinase domain mutations.**

A: Cold treatment of mitotic spindles was performed in T-Rex cells induced to express similar levels of BUBR1- WT, 731X, DKD, and SKD. After endogenous BUBR1 depletion, cells were arrested in mitosis with STLC overnight, followed by release into MG132 for 3.5h. Prior to fixation, the media was removed and replaced with culture media cooled to 4 °C. The dishes were then placed in a shallow ice bath for 10 min before fixation and immunofluorescence staining. Quantitation indicates mitotic spindle staining intensity subtracted from average background measurement.

B: Microtubule intensity measured directly at kinetochores from the experiment in A.

C: Metaphase inter-kinetochore distances measured in cells expressing equalized levels of the BUBR1 pseudokinase domain mutants and aligned at metaphase with MG132 as in Fig 4B. Each dot represents one measurement of inter-KT distance in microns. For each cell analyzed, 5 measurements were made.

D: Hela T-Rex stable cell lines expressing equalized levels of BUBR1 pseudokinase mutants were treated as in Fig 4B. Cells were immunostained with Astrin and CREST to mark kinetochores.

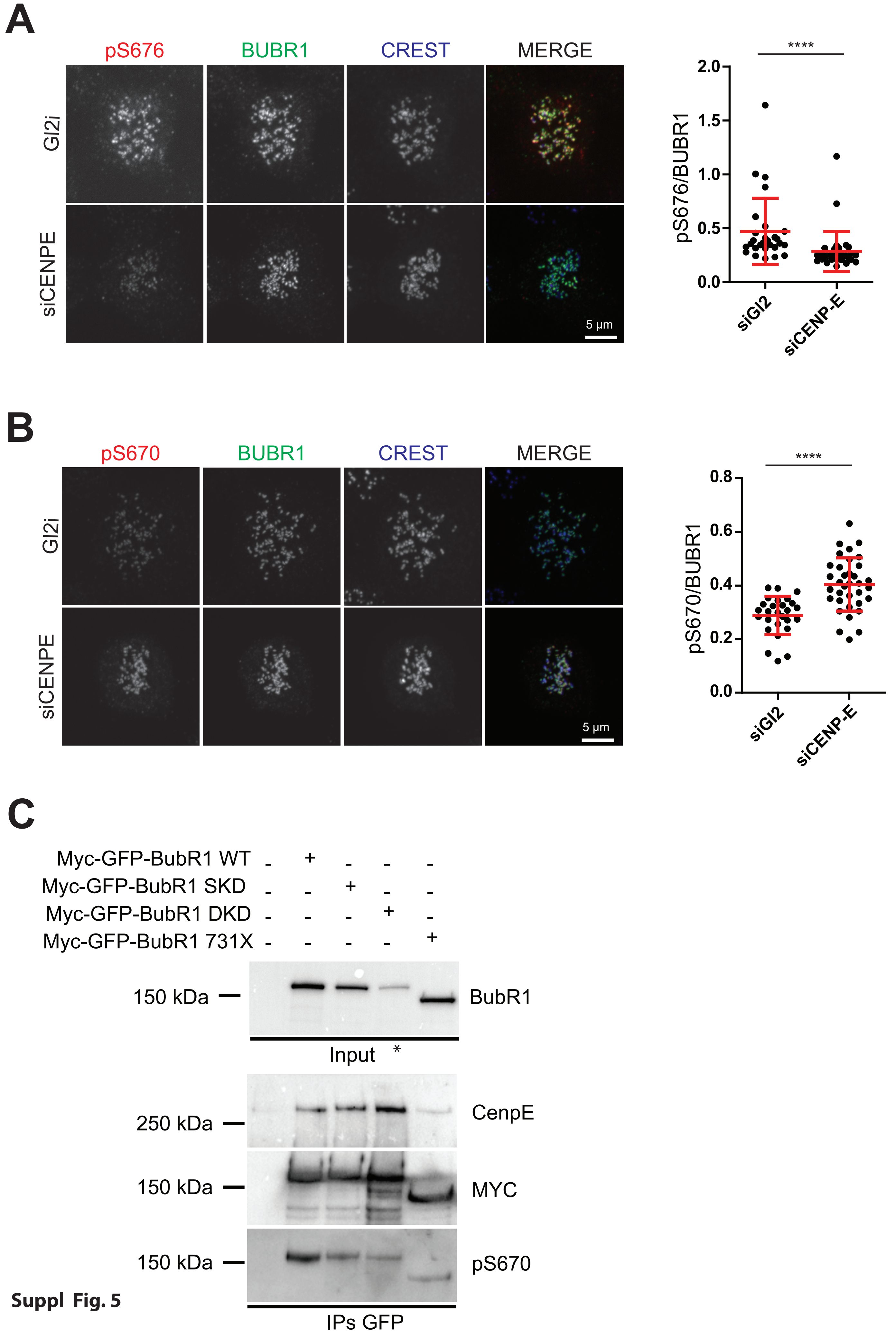
A
pS676
BUBR1
CREST
MERGE
Gl2i
siCENPE
5 µm
****
2.0
1.5
1.0
0.5
0.0
pS676/BUBR1
siGl2
siCENP-E
B
pS670
BUBR1
CREST
MERGE
Gl2i
siCENPE
5 µm
****
0.8
0.6
0.4
0.2
0.0
pS670/BUBR1
siGl2
siCENP-E
C
Myc-GFP-BubR1 WT - + - - -
Myc-GFP-BubR1 SKD - - + - -
Myc-GFP-BubR1 DKD - - - + -
Myc-GFP-BubR1 731X - - - - +
150 kDa
BubR1
Input
*
250 kDa
CenpE
150 kDa
MYC
150 kDa
pS670
IPs GFP

Suppl Fig. 5

**Suppl. Fig 5: Effect of CENP-E depletion on KARD phosphorylation**

A: Hela S3 cells were treated with CENP-E or GL2 siRNAs. Cells were treated with nocodazole after 36h in order to normalize mitotic stages between the two conditions. 48h after transfection, cells were fixed and stained with anti-pS676, BUBR1 and CREST. The quantitation on the right show relative pS676 levels at kinetochores.

B: Cells were treated as in A except immunofluorescence which was performed with Anti-pS670 antibodies, relative quantification of which at kinetochores is shown in the graphic on the right.

C: Cells were transfected with the indicated 3xMYC-GFP-tagged BUBR1 constructs and synchronized in mitosis with nocodazole before lysis and immunoprecipitation with anti-GFP antibodies. * Note that lysates expressing BUBR1-DKD were equalized for the immunoprecipitation experiment. Immunocomplexes were resolved by SDS-PAGE and blotted for CENP-E, pS670 and MYC to demonstrate equal loading.

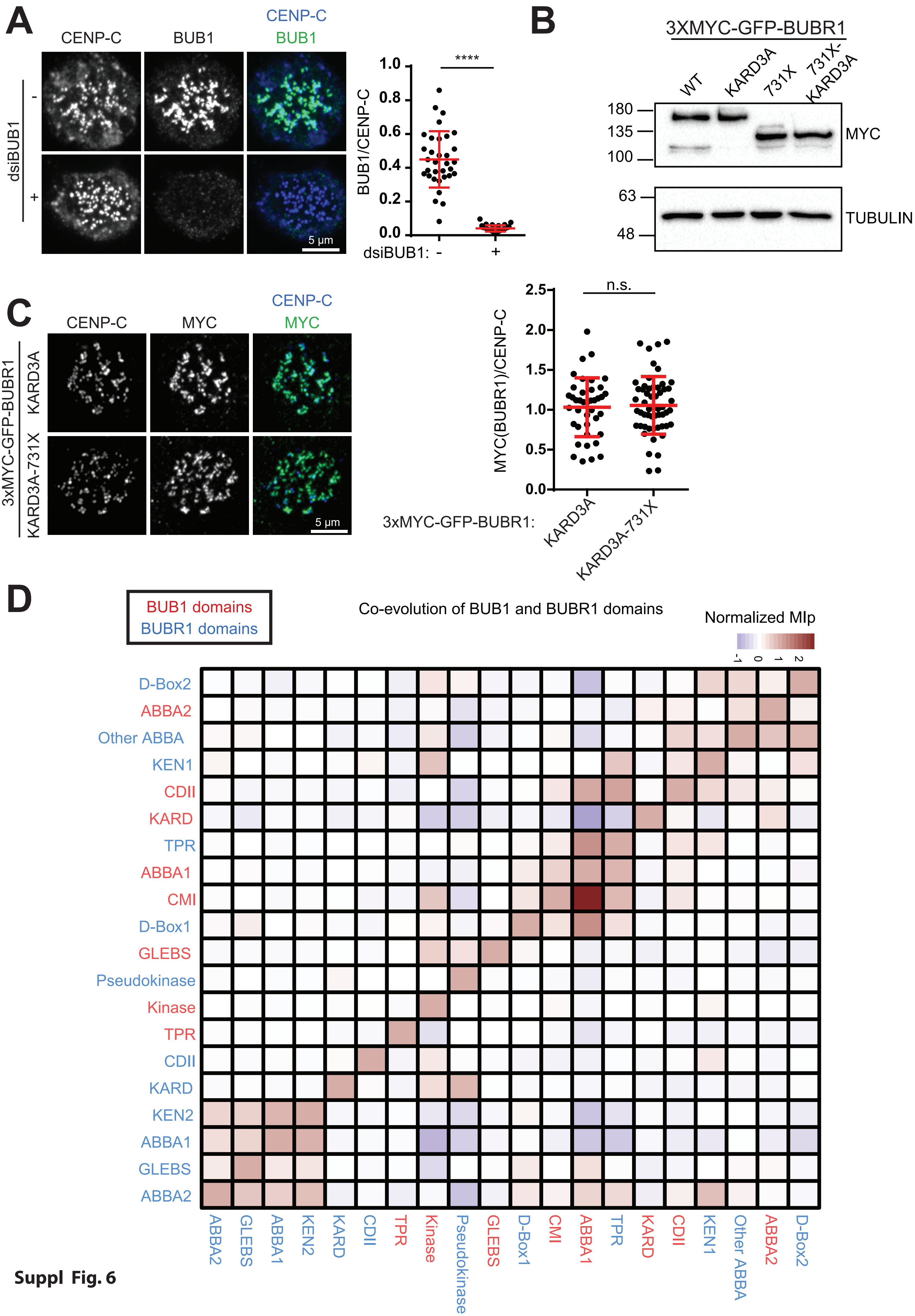
A
CENP-C
BUB1
CENP-C
BUB1
dsiBUB1
-
+
5 µm
BUB1/CENP-C
1.0
0.8
0.6
0.4
0.2
0.0
****
dsiBUB1: -
+
B
3XMYC-GFP-BUBR1
WT
KARD3A
731X
731X-KARD3A
180
135
100
63
48
MYC
TUBULIN
C
CENP-C
MYC
CENP-C
MYC
3xMYC-GFP-BUBR1
KARD3A
KARD3A-731X
5 µm
MYC(BUBR1)/CENP-C
2.5
2.0
1.5
1.0
0.5
0.0
n.s.
3xMYC-GFP-BUBR1:
KARD3A
KARD3A-731X
D
BUB1 domains
BUBR1 domains
Co-evolution of BUB1 and BUBR1 domains
Normalized MIp
-1
0
1
2
D-Box2
ABBA2
Other ABBA
KEN1
CDII
KARD
TPR
ABBA1
CMI
D-Box1
GLEBS
Pseudokinase
Kinase
TPR
CDII
KARD
KEN2
ABBA1
GLEBS
ABBA2
ABBA2
GLEBS
ABBA1
KEN2
KARD
CDII
TPR
Kinase
Pseudokinase
GLEBS
D-Box1
CMI
ABBA1
TPR
KARD
CDII
KEN1
Other ABBA
ABBA2
D-Box2

**Suppl Fig. 6**

**Suppl. Fig 6: BUBR1 and BUB1 domains co-evolution**

A: HeLa cells were treated with GL2 or BUB1 siRNAs for 48 h before fixation and immunofluorescence with BUB1 and CENP-C antibodies. DNA was stained with Hoechst. Quantitation of BUB1 intensity at kinetochores is indicated in the graph on the right.

B: Western blot indicating equalized expression of BUBR1 KARD3A and 731X-KARD3A relative to WT and 731x, respectively.

C: Relative expression levels of BUBR1-KARD3A and BUBR1-KARD3A-731X at kinetochores in nocodazole-arrested stable cell lines.

D: Co-evolution heatmap of BUBR1 and BUB1. The normalized mutual information of a protein (MIp) directly correlates to co-evolving positions in protein families. Domains from BUB1 are shown in red while BUBR1 domains are in blue. The BUBR1 KARD mostly co-evolved with the kinase domain from BUBR1 and BUB1.

**Supplemental Material:**

All sequences, scripts and code use in this study can be viewed at
https://github.com/landrylaboratory/BuBR1_coevolution
